# Vitamin C is an efficient natural product for prevention of SARS-CoV-2 infection by targeting ACE2 in both cell and in vivo mouse models

**DOI:** 10.1101/2022.07.14.499651

**Authors:** Yibo Zuo, Zhijin Zheng, Yingkang Huang, Jiuyi He, Lichao Zang, Tengfei Ren, Xinhua Cao, Ying Miao, Yukang Yuan, Yanli Liu, Feng Ma, Sheng Tian, Jianfeng Dai, Qiang Ding, Hui Zheng

## Abstract

ACE2 is a major receptor for cell entry of SARS-CoV-2. Despite advances in targeting ACE2 to inhibit SARS-CoV-2’s binding, how to efficiently and flexibly control ACE2 levels for prevention of SARS-CoV-2 infection has not been explored. Here, we revealed Vitamin C (VitC) administration as an effective strategy to prevent SARS-CoV-2 infection. VitC reduced ACE2 protein levels in a dose-dependent manner, while partial reduction of ACE2 can greatly restrict SARS-CoV-2 infection. Further studies uncovered that USP50 is a crucial regulator of ACE2 protein levels, and VitC blocks the USP50-ACE2 interaction, thus promoting K48-linked polyubiquitination at Lys788 and degradation of ACE2, without disrupting ACE2 transcriptional expression. Importantly, VitC administration reduced host ACE2 and largely blocked SARS-CoV-2 infection in mice. This study identified an *in vivo* ACE2 balance controlled by both USP50 and an essential nutrient VitC, and revealed a critical role and application of VitC in daily protection from SARS-CoV-2 infection.

**Highlights:**

- VitC reduces ACE2 protein levels in a dose-dependent manner
- VitC and USP50 regulate K48-linked ubiquitination at Lys788 of ACE2
- VitC blocks the interaction between USP50 and ACE2
- VitC administration lowers host ACE2 and prevents SARS-CoV-2 infection *in vivo*

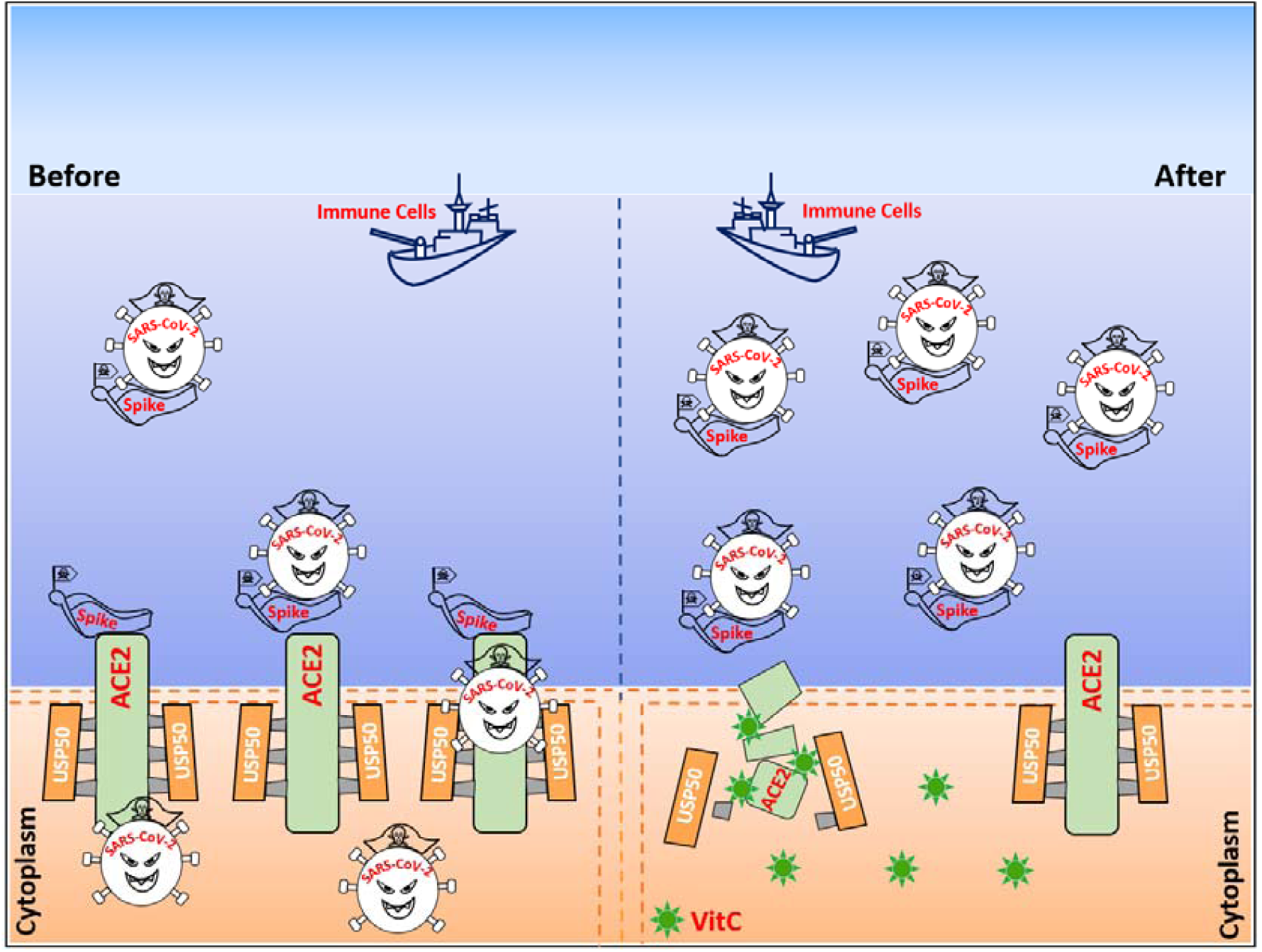

The deubiquitinase USP50 controls ACE2 protein stability and levels, while Vitamin C blocks the USP50-ACE2 interaction and therefore results in ACE2 degradation, offering a flexible and efficient approach to protection of the host from SARS-CoV-2 infection.

## INTRODUCTION

Angiotensin converting enzyme 2 (ACE2) is a regulator of the renin-angiotensin-aldosterone system (RAAS) that maintains blood pressure homeostasis, as well as fluid and salt balance (Donoghue et al., 2000; Krege et al., 1995; Kuba et al., 2006). Recent studies have revealed that ACE2, as a transmembrane glycoprotein, is the major entry receptor of some coronaviruses, including severe acute respiratory syndrome coronavirus 2 (SARS-CoV-2) and SARS-CoV (Hoffmann et al., 2020). Particularly, ACE2 has a 10-20-fold higher affinity to the Spike protein of SARS-CoV-2 than to that of SARS-CoV (Wrapp et al., 2020), which could suggest the severity of SARS-CoV-2-induced coronavirus disease 2019 (COVID-19). In addition, the recently prevalent SARS-CoV-2 mutant Omicron has produced much stronger infectivity, because the Spike trimer of Omicron (BA.2) exhibits 11-fold higher potency in binding to human ACE2 than that of the wild-type (WT) SARS-CoV-2 (Xu et al., 2022). Thus, ACE2 is the key target to control the infection of SARS-CoV-2 and even its current and future mutants.

It has been proved that inhibition of the SARS-CoV-2-Spike recognition of host ACE2 is efficient to block SARS-CoV-2 infection. For this purpose, the current studies have been focused on inhibition of the binding between ACE2 and Spike proteins by two major strategies, including targeting either host ACE2 or the Spike protein of SARS-CoV-2. For example, a small molecule telmisartan can target ACE2 and therefore blocks SARS-CoV-2’s binding (Rothlin et al., 2020). Likewise, many small molecules that target the Spike protein can inhibit ACE2-Spike binding (Wang et al., 2022). In addition, monoclonal antibodies have also been studied to target different proteins for inhibition of ACE2-Spike binding (Chavda et al., 2022). However, these two strategies could be not feasible for timely application to daily prevention of virus infection due to potential harmful side effects from small compounds or antibodies. In addition, ACE2 levels can be upregulated in the elderly by commonly used drugs (Ferrario et al., 2005; Messerli et al., 2018; Verdecchia et al., 2010; Winkelmayer et al., 2005) and also be strongly elevated by SARS-CoV-2 infection (Garvin et al., 2020; Ziegler et al., 2020), which make it difficult to effectively target ACE2.

Based on the consideration for daily protection from SARS-CoV-2 infection, we explored the possibility of partially reducing ACE2 levels as a new strategy for prevention of SARS-CoV-2 infection. Attractively, we noticed that reducing ACE2 protein levels by half can dramatically block SARS-CoV-2 infection. Intriguingly, by screening we found that Vitamin C (VitC, L-ascorbic acid, ascorbate) downregulated ACE2 protein levels in a dose-dependent manner. VitC is an *in vivo* essential nutrient and a natural water-soluble antioxidant that can safely be administered in very large amounts (such as 10 g daily) (Chen et al., 2005). Thus, this study further revealed the mechanisms by which VitC reduces ACE2 protein levels to prevent SARS-CoV-2 infection.

## Results

### VitC effectively reduces ACE2 proteins and restricts cellular infection with SARS-CoV-2

ACE2 expresses in a wide variety of human tissues. For example, ACE2 shows the highest levels in the kidney, small intestine and adipose tissues, the medium levels in the lung, colon and liver tissues, and the lowest levels in the brain, bone marrow and blood (Li et al., 2020). Thus, we observed ACE2 protein levels in several cell lines from different tissues, including HEK293T (kidney), A549 (lung) and Caco-2 (colon), as well as a human fibroblast cell line (2fTGH) due to the reported expression of ACE2 in lung fibroblasts (Mohamed et al., 2021), since these cell lines have been reported to be functional for SARS-CoV-2 infection. We noticed that all these cell lines can more or less express ACE2 proteins (Figure 1A). Given that ACE2 is essential for cell entry of SARS-CoV-2, we first tested how important ACE2 reduction is for protecting the cells from SARS-CoV-2 infection. The results showed that reducing ACE2 levels by only half is enough to tremendously lower the ability of SARS-CoV-2 to enter cells (Figure 1B), suggesting that even partial reduction of ACE2 is able to provide great benefits for the prevention of SARS-CoV-2 infection.

**Figure 1.**
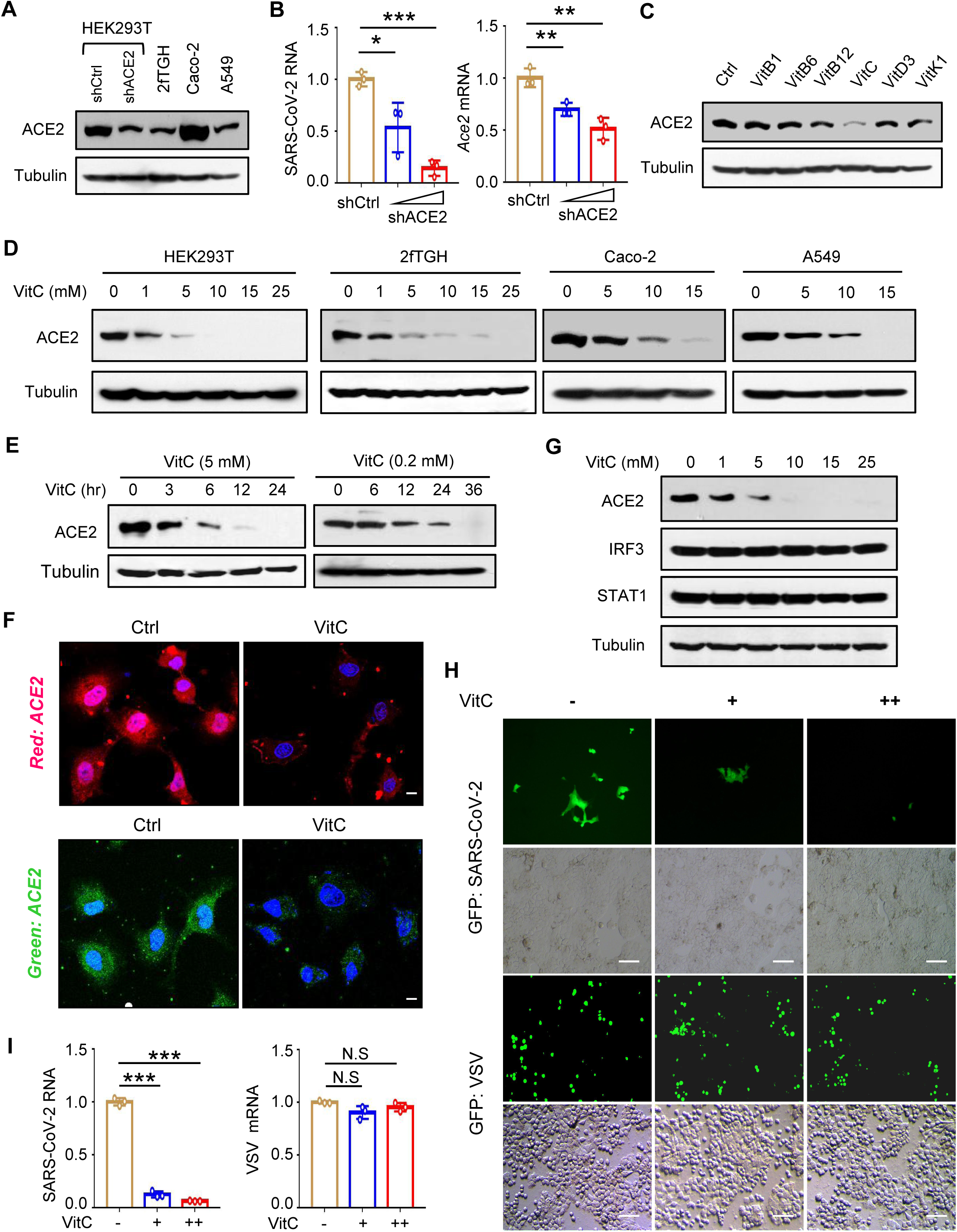
VitC effectively reduces ACE2 proteins and restricts cellular infection with SARS-CoV-2. **(A)** Western blot analysis of endogenous ACE2 in various types of cell lines, including HEK293T transfected with either control shRNAs (shCtrl) or shRNAs against ACE2 (shACE2), 2fTGH, Caco-2 and A549 cells. **(B)** HEK293T cells were transfected with shCtrl or increasing amounts of shACE2. Then cells were subjected to RT-qPCR analysis of *Ace2* mRNA levels (right), or were infected with SARS-CoV-2 GFP/ΔN (MOI = 0.1) for 2 hrs. RT-qPCR was used to analyze SARS-CoV-2 RNA levels (left). **(C)** Western blot analysis of ACE2 in Caco-2 cells treated with vitamin (Vit) compounds (VitB1, 500 µM; VitB6, 500 µM; VitB12, 50 nM; VitC, 5 mM; VitD3, 25 µM; VitK1, 0.5 µM) for 24 hrs. **(D)** Western blot analysis of ACE2 in HEK293T, 2fTGH, Caco-2 and A549 cells treated with VitC at indicated concentrations for 24 hrs. **(E)** Western blot analysis of ACE2 in HEK293T cells treated with 5 mM or 0.2 mM of VitC for different durations. **(F)** Immunofluorescence analysis of ACE2 proteins in HeLa cells treated with VitC (5 mM) for 24 hrs. DAPI was used for the nucleus. Scale bars, 1 μm. **(G)** Western blot analysis of ACE2, IRF3 and STAT1 in A549 cells treated with VitC at indicated concentrations for 24 hrs. **(H)** Fluorescence microscopy of the SARS-CoV-2 GFP/ΔN or VSV-GFP viruses in Caco-2-N cells pretreated with VitC (5 mM and 10 mM) for 24 hrs, and then infected with SARS-CoV-2 GFP/ΔN (MOI = 0.1) or VSV-GFP (MOI = 0.1) viruses for 24 hrs. Scale bar: 100 µm. **(I)** RT-qPCR analysis of SARS-CoV-2 GFP/ΔN or VSV RNA levels in Caco-2 cells pretreated with VitC as (H), and then infected with SARS-CoV-2 GFP/ΔN (MOI = 0.1) or VSV (MOI = 0.1) for 2 hrs. Data are representative of three independent experiments (A, C-F), or are shown as mean and s.d. of three biological replicates (B, I). N.S, not significant, \**p* < 0.05, \*\**p* < 0.01, \*\*\**p* < 0.001 (two-tailed unpaired Student’s *t*-test). See also Figure S1.

Given that new unknown compounds and even known clinical drugs are generally not feasible for timely application to prevention of virus infection, we considered whether some conventional supplements, such as metal elements (Figure S1A) or vitamins (Figure 1C), could effectively downregulate ACE2 and therefore greatly prevent SARS-CoV-2 infection. Interestingly, we found that VitC at a concentration of 5 mM can decrease ACE2 protein levels by around 90% (Figure 1C). It has been reported that normal cells are unaffected by 20 mM VitC (Chen et al., 2005) and even a dose of 49 mM is still well tolerated (Stephenson et al., 2013). Although the concentration of VitC in human plasma is around 50-70 µM under physiological conditions (Lykkesfeldt and Tveden-Nyborg, 2019), the concentrations of VitC in various tissues are dozens-fold higher than that in plasma, in the range of 0.5-10 mM (Lindblad et al., 2013; Lykkesfeldt and Tveden-Nyborg, 2019). When additional VitC is supplemented orally, the plasma concentrations of VitC can raise up to 220 µM (Levine et al., 1996; Lindblad et al., 2013), which could raise the concentrations of VitC in different tissues to much higher than 0.5-10 mM after oral VitC administration. Interestingly, we found that 1-5 mM VitC is enough to strongly reduce ACE2 protein levels in all these cells, particularly in human kidney cells HEK293T and human fibroblasts 2fTGH (Figure 1D). Therefore, we mainly employed these two cell lines in the following studies. In fact, VitC even if at lower concentrations (0.2 mM) can also obviously reduce ACE2 levels in a time-dependent manner (Figure 1E). In addition, by immunofluorescence analysis we further confirmed the reduction of ACE2 proteins by VitC (Figure 1F). Importantly, not all cellular proteins were downregulated by VitC at these concentrations. For example, the protein levels of two commonly studied immune-related transcription factors, IRF3 and STAT1, remained unchanged under 1-25 mM VitC treatment (Figure 1G).

Next, we employed a SARS-CoV-2 GFP/ΔN virus, in which a GFP reporter gene is used to replace the viral nucleocapsid (N) gene in a complete SARS-CoV-2 virus (Ju et al., 2021), to observe the effect of VitC on SARS-CoV-2 infection. To this end, the Caco-2-N cells that stably express the N protein of SARS-CoV-2 were pretreated with VitC, and then infected with the viruses for 24 hrs. We noticed that VitC markedly inhibited infection of SARS-CoV-2 but not the Vesicular Stomatitis Virus (VSV)-GFP virus (Figure 1H; Figure S1B). However, when the cells were first infected with SARS-CoV-2 and then treated with VitC, the SARS-CoV-2 replication in cells was not significantly affected (Figure S1C). Furthermore, the Caco-2-N cells were infected with SARS-CoV-2 GFP/ΔN viruses for only 2 hrs to observe the entry of the viruses. Similarly, we found that VitC treatment strongly blocked cell entry of SARS-CoV-2 but not VSV (Figure 1I). In addition, we also used a SARS-CoV-2-S pseudovirus to study the viral infection mediated by the Spike (S) protein of SARS-CoV-2. Consistently, the results showed that VitC but not either VitB1 or VitD3 dramatically blocked SARS-CoV-2-S pseudovirus infection (Figure S1D). Taken all together, these findings suggested that VitC can greatly block cellular infection with SARS-CoV-2.

### VitC regulates K48-linked polyubiquitination and protein stability of ACE2

We next studied how VitC reduces ACE2 protein levels. Our data showed that VitC did not significantly affect ACE2 mRNA levels (Figure 2A). Thus, we further observed whether VitC regulates ACE2 at the protein level. A cycloheximide (CHX) pulse chase assay demonstrated that VitC treatment promoted ACE2 protein degradation (Figure 2B), suggesting that VitC lowered ACE2 protein stability. Consistent with the above findings, the levels of exogenously expressed Myc-ACE2 proteins were also reduced by VitC (Figure 2C). Furthermore, we determined which pathway ACE2 proteins degrade from. To this end, a proteasome inhibitor (MG132) and a lysosome inhibitor methylamine (MA) were utilized. The results showed that MA blocked VitC-mediated downregulation of ACE2 proteins (Figure 2D), suggesting that VitC promotes ACE2 protein degradation through the lysosome pathway.

**Figure 2.**
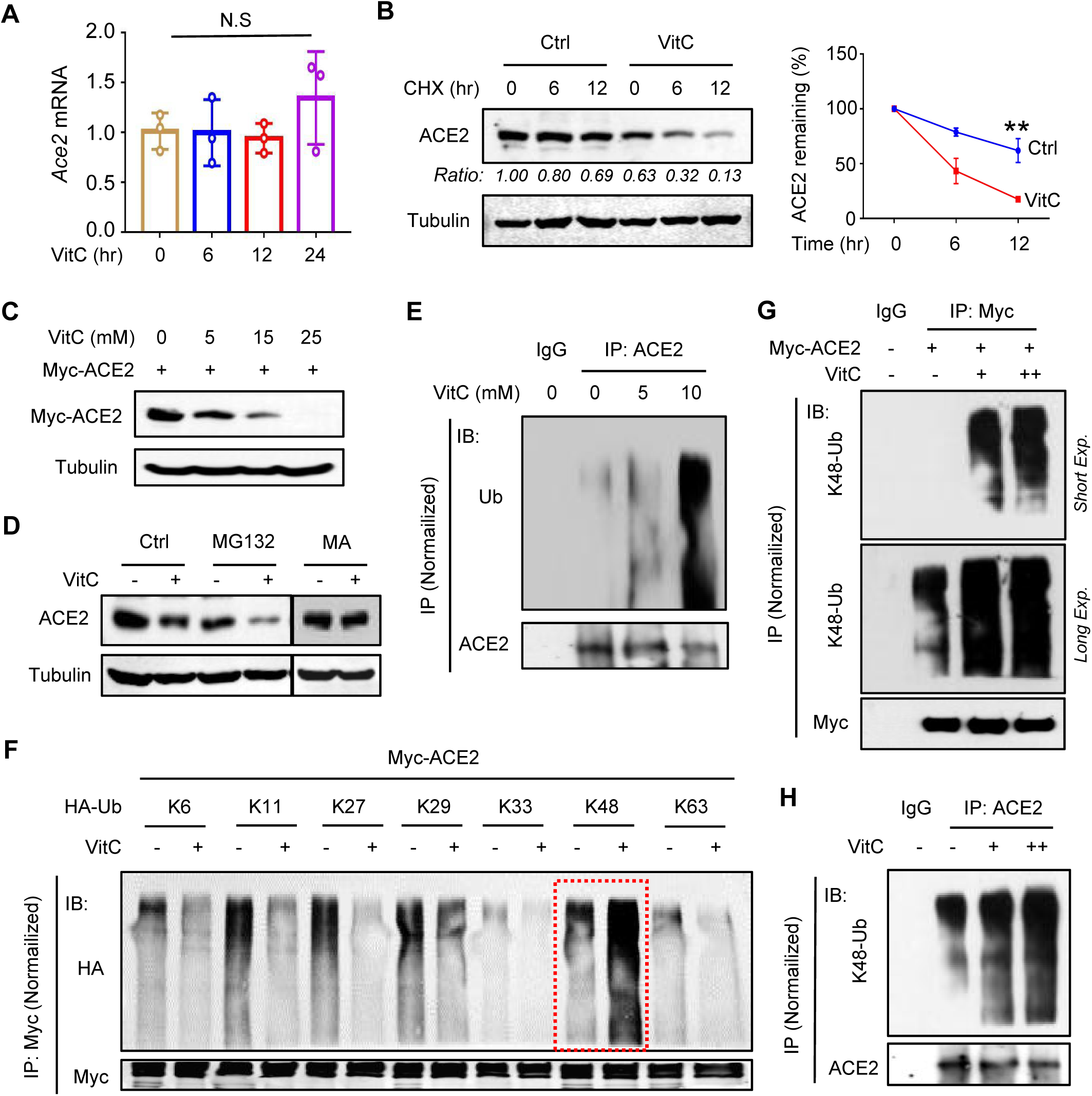
VitC regulates K48-linked polyubiquitination and protein stability of ACE2. **(A)** RT-qPCR analysis of *Ace2* mRNA in 2fTGH cells treated with VitC (5 mM) as indicated. **(B)** Western blot analysis of ACE2 in 2fTGH cells pretreated with ddH_2_O (Ctrl) or VitC (5 mM) for 12 hrs and then treated with CHX (50 μM) for 6 and 12 hrs. **(C)** Western blot analysis of Myc-ACE2 levels in HEK293T cells transfected with Myc-ACE2 and then treated with VitC at indicated concentrations for 12 hrs. **(D)** Western blot analysis of ACE2 in HEK293T cells pretreated with MG132 (10 µM) or MA (10 µM) for 2 hrs, followed by VitC treatment (5 mM) for 6 hrs. **(E)** Immunoprecipitation (IP)-immunoblotting (IB) analysis of ubiquitination (Ub) of endogenous ACE2 in 2fTGH cells treated with VitC at indicated concentrations for 12 hrs. **(F)** IP-IB analysis of ubiquitination types of Myc-ACE2 in HEK293T cells cotransfected with Myc-ACE2 and different types of HA-Ub, and then treated with VitC (5 mM) for 12 hrs. **(G)** IP-IB analysis of K48-linked polyubiquitination (K48-Ub) of Myc-ACE2 in HEK293T cells transfected with Myc-ACE2 and then treated with VitC (2.5 mM and 5 mM) for 12 hrs, using a specific anti-K48-Ub antibody. **(H)** IP-IB analysis of K48-Ub of endogenous ACE2 in 2fTGH cells treated with VitC (2.5 mM and 5 mM) for 12 hrs. Data are representative of three independent experiments (B-H), or are shown as mean and s.d. of three biological replicates (A, B). N.S, not significant, \*\**p* < 0.01 (two-tailed unpaired Student’s *t*-test). See also Figure S2.

Given that VitC promotes ACE2 protein degradation, we next analyzed whether VitC regulates ACE2 ubiquitination. The results showed that VitC treatment remarkably increased the ubiquitination levels of both exogenously expressed ACE2 (Figure S2A) and endogenous ACE2 (Figure 2E; Figure S2B). Furthermore, the analysis of ubiquitination types of ACE2 revealed that in comparison to other types of polyubiquitination linkage, VitC largely promoted K48-linked polyubiquitination of ACE2 (Figure 2F), which was in line with increased degradation of ACE2 proteins, since K48-linked polyubiquitination has been demonstrated to mainly induce protein degradation. Moreover, using a specific anti-K48-ubiquitination antibody, we confirmed that VitC markedly promoted K48-linked polyubiquitination of exogenously expressed ACE2 (Figure 2G) and endogenous ACE2 in multiple types of cells, including HEK293T (Figure 2H), A549 and Caco-2 cells (Figure S2C and Figure S2D). Collectively, these findings revealed that VitC reduces ACE2 protein levels by inducing K48-linked polyubiquitination and lysosome-dependent degradation of ACE2 proteins.

### VitC reduces ACE2 protein levels largely dependently on the deubiquitinase USP50

To identify the key molecule that mediates ACE2 ubiquitination induced by VitC, we fist utilized a pan-deubiquitinase inhibitor, PR619 (Ritorto et al., 2014). The results showed that inhibition of the deubiquitinases substantially decreased ACE2 proteins and blocked VitC-mediated downregulation of ACE2 (Figure 3A), suggesting that VitC could target certain deubiquitinases to reduce ACE2 proteins. Thus, a deubiquitinase-expressing library was used to identify the potential deubiquitinase. We noticed that despite multiple deubiquitinases that could be involved in regulation of ACE2 protein levels, the deubiquitinase USP50 showed the strongest effect on ACE2 upregulation (Figure 3B). Further analysis demonstrated that USP50 was able to interact with exogenously expressed ACE2 (Figure 3C), and actually there was a constitutive interaction between endogenous USP50 and ACE2 in cells (Figure 3D; Figure S3A). Moreover, USP50 overexpression upregulated ACE2 protein levels in a dose-dependent manner (Figure 3E), while knockdown of USP50 by two specific shRNAs dramatically reduced ACE2 levels (Figure 3F). Consistently, USP50 overexpression increased the protein stability of ACE2 (Figure 3G). The results from USP50 knockout cells confirmed that USP50 deficiency reduced ACE2 protein levels (Figure 3H). Importantly, in USP50-deficient cells VitC lost the efficiency to downregulate ACE2 levels (Figure 3I), suggesting that USP50 is critical to VitC-induced ACE2 reduction.

**Figure 3.**
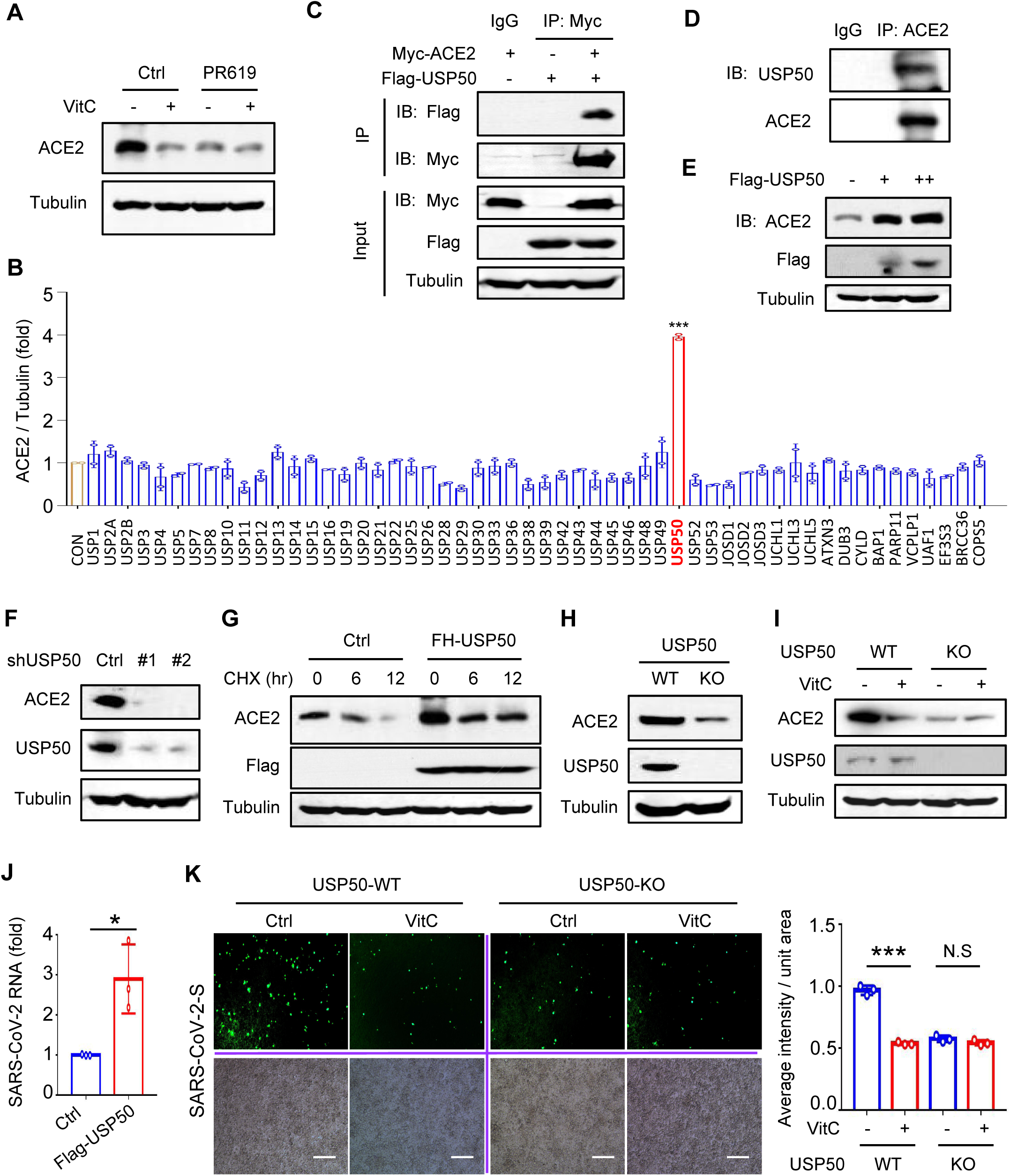
VitC reduces ACE2 levels largely dependently on the deubiquitinase USP50. **(A)** Western blot analysis of ACE2 in HEK293T cells pretreated with PR619 (50 µM, 2 hrs) and then treated with VitC (5 mM) for 12 hrs. **(B)** HEK293T cells were individually transfected with the plasmids from the human DUBs expression library. Western blot was used to identify the key deubiquitinase that significantly increases ACE2 levels. **(C)** IP-IB analysis of the interaction between Flag-USP50 and Myc-ACE2 in HEK293T cells cotransfected with these two constructs. **(D)** Immunoprecipitation analysis of the interaction between endogenous USP50 and ACE2 in 2fTGH cells. **(E)** Western blot analysis of ACE2 in HEK293T cells transfected with increasing amounts of Flag-USP50. **(F)** Western blot analysis of ACE2 in HEK293T cells transfected with shCtrl or shUSP50 (#1 or #2). **(G)** Western blot analysis of ACE2 in HEK293T cells transfected with Flag-USP50 and then treated with CHX (50 μM) as indicated. **(H)** Western blot analysis of ACE2 in *Usp50*^*+/+*^ and *Usp50*^*-/-*^ HEK293T cells. **(I)** Western blot analysis of ACE2 in *Usp50*^*+/+*^ and *Usp50*^*-/-*^ HEK293T cells treated with VitC (5 mM) for 12 hrs. **(J)** RT-qPCR analysis of SARS-CoV-2 GFP/ΔN RNA levels in HEK293T cells transfected with Flag-USP50 and then infected with the SARS-CoV-2 GFP/ΔN virus (MOI = 0.1) for 24 hrs. **(K)** Fluorescence microscopy of the SARS-CoV-2-S pseudovirus in *Usp50*^*+/+*^ and *Usp50*^*-/-*^ HEK293T cells pretreated with or without VitC (5 mM) for 12 hrs, followed by infection with SARS-CoV-2-S pseudovirus (MOI = 0.1) for 24 hrs. Scale bar: 100 µm. Data are representative of three independent experiments (A-I), or are shown as mean and s.d. of three biological replicates (J, K). N.S, not significant, \**p* < 0.05, \*\*\**p* < 0.001 (two-tailed unpaired Student’s *t*-test). See also Figure S3.

Given the importance of USP50 in regulating ACE2 levels by VitC, we further observed the role of USP50 in SARS-CoV-2 infection. Consistent with USP50-mediated ACE2 upregulation, overexpression of USP50 significantly promoted cellular infection with SARS-CoV-2 (Figure 3J). Conversely, knockdown of USP50 inhibited SARS-CoV-2 infection (Figure S3B). Importantly, USP50 knockout largely restricted the effect of VitC on inhibition of SARS-CoV-2 infection (Figure 3K), suggesting that USP50 is crucial for VitC to inhibit SARS-CoV-2 infection. Taken together, these findings demonstrated that VitC regulates ACE2 levels and SARS-CoV-2 infection dependently on USP50.

### VitC and USP50 regulate K48-linked polyubiquitination at Lys788 of ACE2

Next, we sought to determine the detailed mechanisms by which the deubiquitinase USP50 regulates ACE2. We first found that a deubiquitinase inactive mutant of USP50 (USP50-C53S) lost the ability to upregulate ACE2 protein levels (Figure S3C), suggesting that USP50 regulates ACE2 levels dependently on its deubiquitinase activity. In line with this finding, USP50-wild type (WT) but not the USP50-C53S mutant reduced the ubiquitination levels of ACE2 (Figure 4A; Figure S3D), while USP50 knockdown increased ACE2 ubiquitination (Figure 4B). The analysis of ubiquitination types of ACE2 revealed that USP50 mainly reduced K48-linked polyubiquitination of ACE2, compared to other types of ubiquitination linkage (Figure 4C), which is in accord with USP50-mediated upregulation of ACE2 protein levels. Furthermore, using a K48-linked ubiquitination antibody, we confirmed that USP50 regulates K48-linked polyubiquitination of ACE2 (Figure 4D).

**Figure 4.**
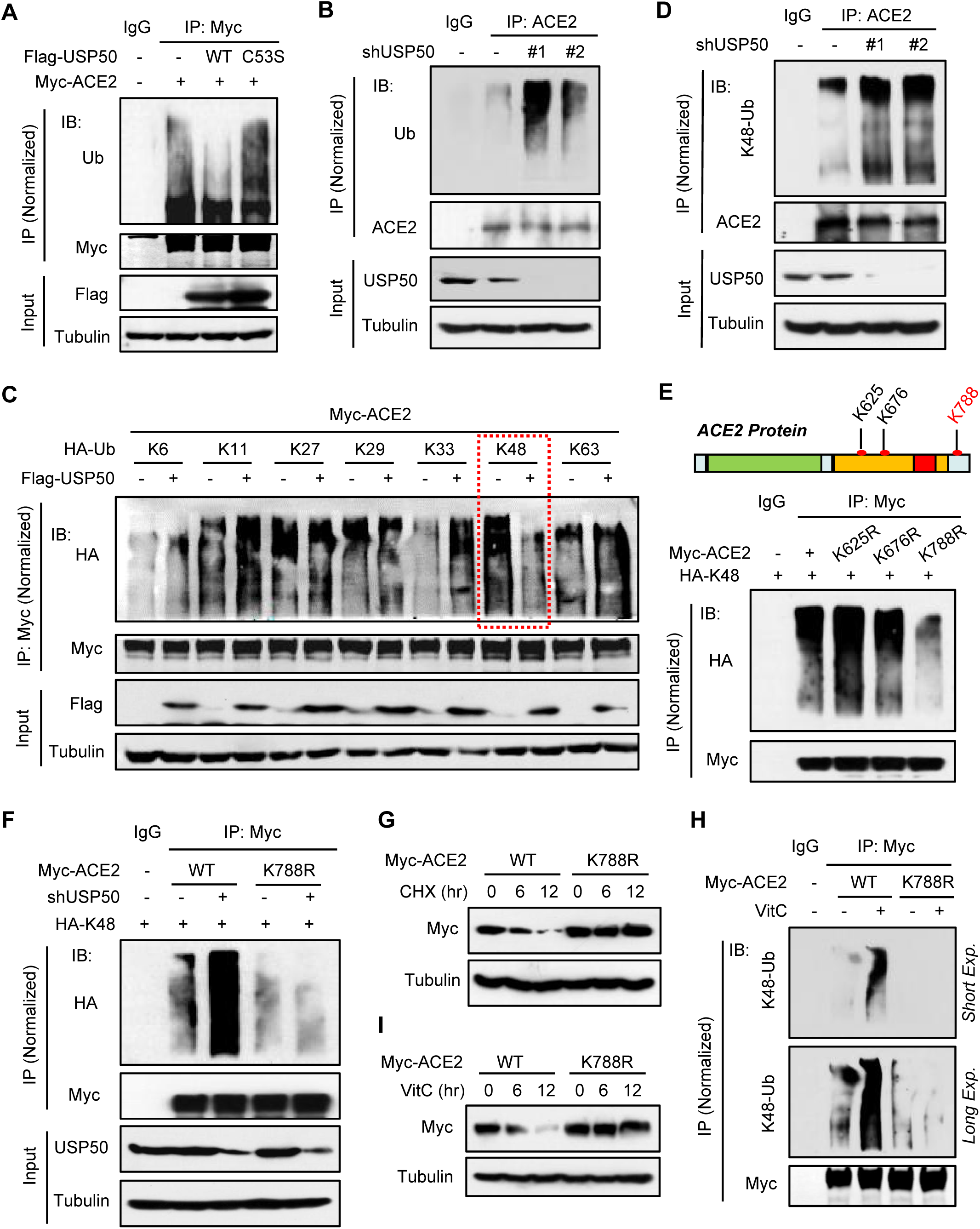
USP50 regulates K48-linked polyubiquitination at Lys788 of ACE2. **(A)** IP-IB analysis of ubiquitination of Myc-ACE2 in HEK293T cells transfected with Myc-ACE2, together with Flag-USP50 (WT) or its deubiquitinase inactive mutant (C53S). **(B)** IP-IB analysis of ubiquitination of endogenous ACE2 in HEK293T cells transfected with either shCtrl (-) or shUSP50 (#1, #2). **(C)** IP-IB analysis of ubiquitination types of Myc-ACE2 in HEK293T cells cotransfected with Myc-ACE2, Flag-USP50 and different types of HA-Ub. **(D)** IP-IB analysis of K48-Ub of endogenous ACE2 in HEK293T cells transfected with shCtrl (-) or shUSP50 (#1, #2). **(E)** Putative ubiquitination sites of ACE2 in the PhosphoSitePlus database (Upper). Myc-ACE2 K48-linked ubiquitination was analyzed by IP-IB in HEK293T cells cotransfected with Myc-ACE2 (WT or its mutants) and HA-K48-Ub (Lower). **(F)** IP-IB analysis of Myc-ACE2 K48-linked ubiquitination in HEK293T cells cotransfected with Myc-ACE2 (WT or K788R) and HA-K48, together with shCtrl or shUSP50. **(G)** Western blot analysis of Myc-ACE2 in HEK293T cells transfected with Myc-ACE2 (WT or K788R) and then treated with CHX (50 μM) as indicated. **(H)** IP-IB analysis of K48-Ub of Myc-ACE2 in HEK293T cells transfected with Myc-ACE2 (WT or K788R) and then treated with VitC (5 mM) for 12 hrs, by a specific anti-K48-Ub antibody. **(I)** Western blot analysis of Myc-ACE2 in HEK293T cells transfected with Myc-ACE2 (WT or K788R) and then treated with VitC (5 mM) as indicated. Data are representative of three independent experiments (A-I). See also Figure S3 and S4.

We further explored the key lysine (Lys) residue of ACE2 regulated by USP50. From the PhosphoSitePlus database, we noticed that there are three ubiquitinated lysine residues on ACE2, including Lys 625, 676, and 788 (Figure 4E). Then, each lysine (K) residue was mutated to arginine (R). We noticed that mutation of the Lys788 residue largely lowered the K48-linked polyubiquitination levels of ACE2 (Figure 4E; Figure S4A). Furthermore, when the Lys788 residue of ACE2 was mutated, USP50 knockdown cannot regulate both K48-linked polyubiquitination (Figure 4F) and protein levels (Figure S4B) of ACE2 any longer, suggesting that USP50 regulates K48-linked polyubiquitination at the Lys788 residue of ACE2. In addition, mutation of the Lys788 residue increased the protein stability of ACE2 (Figure 4G). Importantly, VitC-induced upregulation of ACE2 K48-linked polyubiquitination was abolished by mutating the Lys788 residue of ACE2 (Figure 4H). Consistently, VitC cannot reduce the protein levels of ACE2-K788R (Figure 4I). Taken all together, these findings demonstrated that VitC and USP50 regulate K48-linked polyubiquitination at Lys 788 of ACE2.

### VitC blocks the interaction between USP50 and ACE2

Given that VitC has a similar effect as USP50 knockdown in regulating ACE2 ubiquitination, we hypothesized that VitC could inhibit USP50-mediated regulation of ACE2. Thus, we employed an *in vitro* binding assay to observe the potential interaction of VitC with USP50 or ACE2 (Figure 5A). The results showed that Myc-ACE2 but not Flag-USP50 proteins can significantly bind with VitC (Figure 5B), suggesting VitC could interact with ACE2. Furthermore, we used a recombinant human ACE2 protein to analyze the binding of VitC to the pure ACE2 proteins. The results confirmed the interaction between VitC and pure ACE2 (Figure 5B). Next, our data demonstrated that the amino acid 201-400 domain of ACE2 is critical for binding of VitC (Figure 5C). Thus, we further evaluated their binding conformation. Based on the Glide docking module in the Schrödinger molecular simulation software package, we used standard precision (SP) to dock VitC to the binding pocket of ACE2 (Figure 5D). The docking diagram indicates that multiple amino acids of ACE2 was involved in the interaction with VitC (Figure 5E).

**Figure 5.**
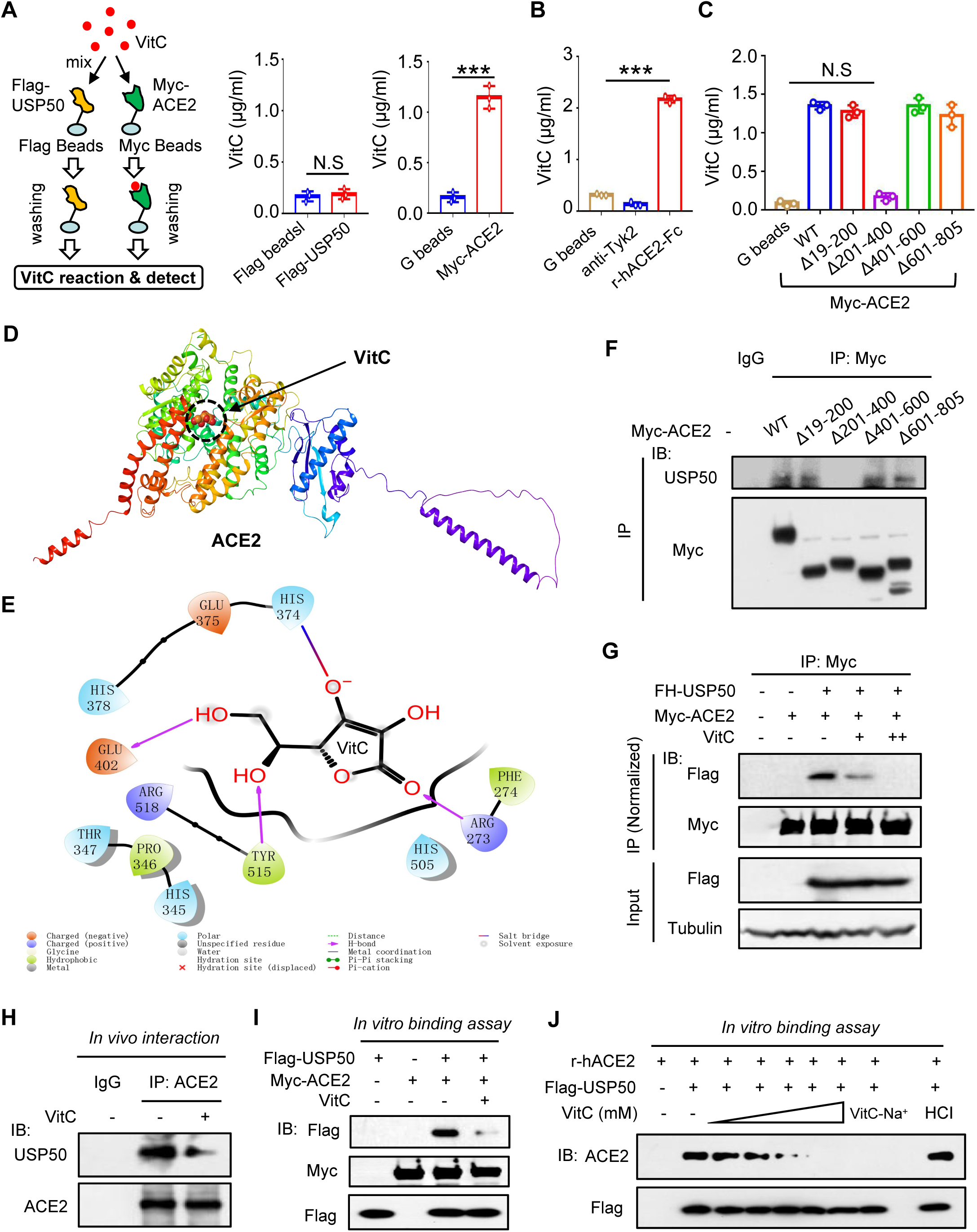
VitC blocks the interaction between USP50 and ACE2. **(A)** The procedure for analysis of the binding of VitC to Flag-USP50 or Myc-ACE2 (left) can be seen detailedly in the Methods. The concentrations of VitC that binds with either Flag-USP50 (middle) or Myc-ACE2 (right) proteins were measured by the Vitamin-C Detection Kit. **(B)** Recombinant human ACE2-IgG-Fc proteins (r-hACE2-Fc) or anti-Tyk2 IgG proteins were incubated with protein-G beads for 2 hrs, and then VitC was added for binding. After washing and centrifuging, the concentrations of VitC that binds to anti-Tyk2 proteins or r-hACE2-Fc proteins were detected as (A). **(C)** The concentrations of VitC that binds to Myc-ACE2 (WT) or its deletion mutants were detected as (A). **(D)** VitC docking to the binding pocket of ACE2 by standard precision (SP) using the Glide docking module in the Schrödinger molecular simulation software. **(E)** The docking diagram between amino acids of ACE2 and VitC based on the SP docking scoring function. **(F)** IP-IB analysis of the interaction between USP50 and Myc-ACE2 (WT) or its deletion mutants (Δ19-200, Δ201-400, Δ401-600 and Δ601-805) in HEK293T. **(G)** IP-IB analysis of the interaction between Flag-USP50 and Myc-ACE2 in HEK293T cells transfected with these two constructs and then treated with VitC (2.5 mM and 5 mM) for 12 hrs. **(H)** IP-IB analysis of the *in vivo* interaction between endogenous USP50 and ACE2 in 2fTGH cells treated with VitC (5 mM) for 12 hrs. **(I)** Myc-ACE2 and Flag-USP50 proteins were immunoprecipitated from HEK293T cells transfected with either Myc-ACE2 or Flag-USP50. Flag-USP50 proteins were eluted by the Flag (M2) agarose. After washing, Flag-USP50 proteins were mixed with the Myc beads with Myc-ACE2, together with or without VitC (5 mM) for 2 hrs. After centrifuging, Flag-USP50 proteins interacting with Myc-ACE2 were analyzed by immunoblotting. **(J)** Flag-USP50 proteins were obtained as (I). Flag-USP50 and r-hACE2 proteins were mixed, together with increasing amounts of VitC or with VitC-Na^+^ (20 mM). HCl is a pH control (pH = 4). After 2 hrs incubation, ACE2 proteins were analyzed by immunoblotting by a specific anti-ACE2 antibody. Data are representative of three independent experiments (F-J), or are shown as mean and s.d. of three biological replicates (A-C). N.S, not significant. \*\*\**p* < 0.001 (two-tailed unpaired Student’s *t*-test). See also Figure S5.

Interestingly, further analysis revealed that the amino acid 201-400 domain of ACE2 is also critical for the interaction of USP50 with ACE2 (Figure 5F), suggesting that VitC could compete with USP50 to bind with ACE2. In line with the speculation, VitC inhibited the interaction between Flag-USP50 and Myc-ACE2 in cells in a dose-dependent manner (Figure 5G). In fact, VitC did not affect USP50 protein levels (Figure S5). Similarly, VitC can also block the interaction between endogenous USP50 and ACE2 (Figure 5H). An *in vitro* binding assay demonstrated the competition between VitC and USP50 to interact with ACE2 (Figure 5I). Moreover, using a recombinant human ACE2, we confirmed that VitC can block the interaction between USP50 and ACE2 in a dose-dependent manner (Figure 5J). Collectively, these findings suggested that VitC competitively inhibits the USP50-ACE2 interaction.

### VitC administration reduces ACE2 and restricts SARS-CoV-2 infection *in vivo*

To observe how VitC administration benefits prevention of SARS-CoV-2 infection *in vivo*, we employed a humanized ACE2 (hACE2) mouse model, since the Spike protein of SARS-CoV-2 cannot target mouse ACE2. We noticed that USP50 indeed interacts with ACE2 constitutively in all observed tissues, including the lung, liver and kidney (Figure 6A; Figure S6A). It has been reported that intraperitoneal (*i*.*p*.) administration of VitC (1 g/kg of body weight) in mice can give maximum plasma concentration of 7 mM (Vilcheze et al., 2018). Thus, based on the used concentration of VitC (1-5 mM) in cell lines in this study, we observed the *in vivo* effects of VitC administration (0.3 g/kg of body weight) on ubiquitination and protein levels of ACE2, as well as SARS-CoV-2-S infection. The results showed that VitC strongly blocked the interaction between USP50 and ACE2 in lung tissues of mice (Figure 6B). Consistently, the K48-linked polyubiquitination levels of endogenous ACE2 in mouse tissues were remarkably upregulated by VitC administration (Figure 6C). As a consequence, VitC administration largely reduced the levels of ACE2 proteins in different tissues of mice (Figure 6D and E; Figure S6B).

**Figure 6.**
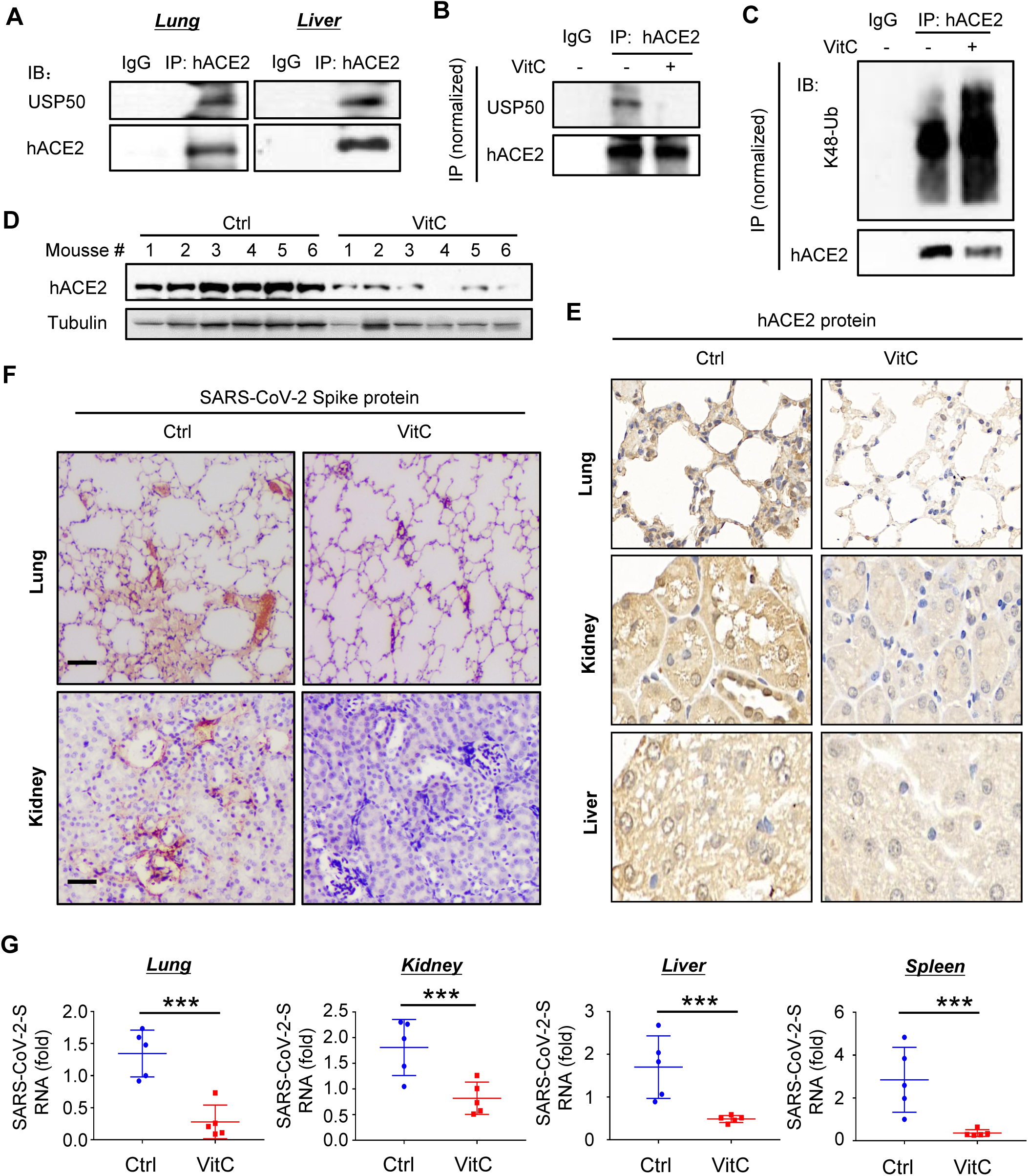
VitC administration reduces ACE2 and restricts SARS-CoV-2 infection *in vivo*. **(A)** IP-IB analysis of the interaction between endogenous hACE2 and USP50 in the lung and liver tissues of hACE2 mice. **(B)** The hACE2 mice were intraperitoneally administrated with VitC (300 mg/day/kg body weight) for two days. The interaction between USP50 and hACE2 in mouse lung tissues was analyzed by IP-IB. **(C)** IP-IB analysis of K48-Ub of hACE2 in mouse lung tissues from (B). **(D)** Western blot analysis of hACE2 levels in lung tissues of hACE2 mice administrated with VitC as (B). **(E)** Immunohistochemical staining of hACE2 protein in the lung, kidney and liver tissues from (B). **(F)** The hACE2 mice were administrated with VitC as (B). Mice were then given intraperitoneal injections of SARS-CoV-2-S pseudoviruses (1×10^6^ PFU per gram body). After 24 hrs, immunohistochemical staining was performed to analyze the SARS-CoV-2 Spike proteins in mouse lung and kidney tissues. Scale bar: 100 µm. **(G)** RT-qPCR analysis of the SARS-CoV-2 Spike mRNA levels in lung, kidney, liver and spleen tissues of hACE2 mice treated with VitC and SARS-CoV-2-S pseudoviruses as (F). Data are representative of three independent experiments (A-D). All graphs show the mean ± SEM for five individual mice (G). \*\*\**p* < 0.001 (two-tailed unpaired Student’s *t*-test). See also Figure S6 and S7.

Given that VitC administration substantially lowers ACE2 levels, we further utilized a SARS-CoV-2-S pseudovirus to study the SARS-CoV-2 Spike-mediated infection of the hACE2 mice. To this end, the mice were first administrated with VitC (0.3 g/day/kg of body weight) for two days and then were infected with the SARS-CoV-2-S viruses for 24 hrs. By immunostaining the Spike proteins of SARS-CoV-2, we found that VitC markedly lowered the levels of SARS-CoV-2 Spike proteins in various mouse tissues (Figure 6F), suggesting that VitC inhibits infection of the SARS-CoV-2-S viruses *in vivo*. Moreover, the analysis of the SARS-CoV-2-S RNA levels confirmed that VitC administration significantly restricted SARS-CoV-2-S virus infection (Figure 6G). Taken all together, these findings demonstrated that VitC administration effectively blocks SARS-CoV-2 infection by reducing ACE2 levels *in vivo*.

## DISCUSSION

VitC administration is a flexible and easy way to be used according to daily actual requirements. This study revealed that VitC, as a natural compound from a wide range of sources, can efficiently reduce ACE2 protein levels in a dose-dependent manner, thus providing great benefits to prevention of SARS-CoV-2 infection (Figure S7). Besides a role in mediating SARS-CoV-2 cell entry, ACE2 also has some important physiological functions. ACE2 plays a crucial role in the cardiovascular system by regulating the RAAS (Crackower et al., 2002; Donoghue et al., 2000). Activation of ACE2-Ang I (1-7) axis regulates inflammatory responses and protects against organ injury in many diseases, such as cardiovascular disease, chronic kidney disease, obesity, liver and lung injury (Rodrigues Prestes et al., 2017). It’s well known that in addition to respiratory symptoms, COVID-19 also results in some extrapulmonary pathologies, including vasculature and myocardial complications, kidney injury, and hepatic injury, because ACE2 is widely expressed in many tissues (Gupta et al., 2020). These suggest that ACE2 is actually important for protection from tissue injury at the late stage of COVID-19. Thus, this study provided evidence that VitC could be not good for the therapy of COVID-19 at the late stage of SARS-CoV-2 infection, which many clinical therapeutic studies actually have been performing for COVID-19, since VitC dramatically reduces ACE2 levels. Here, our study suggested that VitC administration is an efficient strategy for daily protection from SARS-CoV-2 infection, and also possibly for the therapy of early SARS-CoV-2 infection.

VitC has highly differential distribution in the body. Under physiological conditions, the concentration of VitC in the plasma of healthy individuals is 50-80 µM (Lykkesfeldt and Tveden-Nyborg, 2019). However, VitC concentrations in various tissues are much higher than that in the plasma, ranging from 0.2-10 mM under physiological conditions: 2-10 mM in the brain and adrenal, about 1mM in the lung and liver, 0.3-0.5 mM in the kidney (Lindblad et al., 2013; Lykkesfeldt and Tveden-Nyborg, 2019). When administrating with VitC, it could result in great improvement in VitC concentrations in both plasma and tissues. VitC has very big tolerated doses *in vivo*. It has been reported that the tolerated oral dose of 3 g every 4 hours and an intravenous dose of 50 g were predicted to give the peak plasma VitC concentrations of 220 µM and 13,400 µM, respectively (Padayatty et al., 2004). And a dose of VitC (2 g, three time/day) could result in a steady plasma concentration of around 250 µM (Lykkesfeldt and Tveden-Nyborg, 2019; Nielsen et al., 2015). Our study demonstrated that VitC at a concentration of 1-10 mM in both cell models and a mouse *in vivo* model, which is close to the physiological concentrations of VitC in tissues and at least can be easily achieved under VitC oral administration, lowered ACE2 protein levels in a dose-dependent manner. Thus, we believe that sufficient VitC-containing daily diets are necessary for maintaining efficient VitC concentrations in tissues to restrict SARS-CoV-2 infection, while additional VitC oral administration can provide greater benefits to the protection of the body from SARS-CoV-2 infection. Importantly, this study demonstrated that VitC did not affect normal transcriptional expression of ACE2, suggesting that ACE2 expression is not irreversibly disrupted in cells and will be recovered when VitC administration stops or is restricted to lower dosages.

The elderly are the most serious victims of COVID-19. SARS-CoV-2 is highly infective with considerable fatality rate in the elderly (Uhler and Shivashankar, 2020). In fact, ACE2 levels often have a significant increase in the elderly. For elderly patients with various diseases, such as refractory hypertension, coronary artery disease, and heart failure, the angiotensin converting enzyme inhibitors (ACEIs) and angiotensin II receptor blockers (ARBs) are highly recommended for the management of cardiovascular diseases (Messerli et al., 2018; Verdecchia et al., 2010), as well as diabetes and renal insufficiency (Winkelmayer et al., 2005). However, ACEIs and ARBs can increase the numbers of ACE2 receptors in the cardiopulmonary circulation (Ferrario et al., 2005). In addition, during RAAS overactivation, ACE2 levels can be upregulated, which could promote SARS-CoV-2 cell entry (Edenfield and Easley, 2022). It has been reported that patients with COVID-19 infections, and most likely treated with ACEIs or ARBs, suffered more severe disease outcomes (Guan et al., 2020). Interestingly, ACE2 is an interferon (IFN)-stimulated gene (ISG) (Ziegler et al., 2020) Thus, Elevated IFN during SARS-CoV-2 infection increases ACE2 expression in cells. ACE2 levels have been reported to increase 199-fold in cells in bronchoalveolar lavage fluid (BALF) from COVID-19 patients (Garvin et al., 2020). Thus, reducing excessive ACE2 is necessary to lower the risk of SARS-CoV-2 infection and alleviate the severity of COVID-19, in particular for the elderly.

Despite the importance of ACE2 in regulating RAAS and SARS-CoV-2 cell entry, the regulation of ACE2 ubiquitination and protein stability remains largely unexplored. The E3 ubiquitin ligase MDM2 (Shen et al., 2020) and Skp2 (Wang et al., 2021) were recently reported to induce ACE2 ubiquitination. However, how the deubiquitinases control ACE2 ubiquitination and protein levels is unknown. Our study for the first time identified the deubiquitinase USP50 as a crucial regulator of ACE2 protein levels in cells. Knockdown of USP50 or inhibition of the USP50 activity extremely reduced cellular levels of ACE2 protein (Figure 3A and 3F). Recent studies showed that USP50 is involved in fatty acid oxidation (Li et al., 2022) and erythropoiesis (Cai et al., 2018). Our finding suggested the significances of USP50 in controlling both SARS-CoV-2 infection and regulating RAAS, which could suggest a new target for the therapy of certain viral infections and cardiovascular diseases. In summary, this study establishes a link between an essential nutrient VitC and cellular USP50-ACE2 regulation, and could provide an easy but efficient strategy for daily protection of the body from SARS-CoV-2 infection.

## STAR METHODS

Detailed methods are provided in the online version of this paper and include the following:

- KEY RESOURCES TABLE
- CONTACT FOR REAGENT AND RESOURCE SHARING
- EXPERIMENTAL MODEL AND SUBJECT DETAILS
  - Mice
- METHOD DETAILS
  - Cell culture and reagents
  - Plasmids and transfection
  - Liver, lung and kidney histology
  - VitC detection assay
  - *In vitro* protein-protein binding assay
  - CRISPR-Cas9 genome editing
  - VitC docking with ACE2
  - Immunoblotting (IB) and Immunoprecipitation (IP)
  - RNA isolation and quantitative real-time PCR
  - Virus and viral infection *in vitro*
  - Viral infection *in vivo*
  - Immunofluorescence microscopy
- QUANTIFICATION AND STATISTICAL ANALYSIS
  - Statistical analysis
- DATA AND SOFTWARE AVAILABILITY

## STAR METHODS

### KEY RESOURCES TABLE

| REAGENT or RESOURCE | SOURCE | IDENTIFIER |
| --- | --- | --- |
| Antibodies |  |  |
| Anti-ACE2 | Affinity | Cat.# AF5165<br>RRID: AB_2837651 |
| Anti-human ACE2 | Cell Signaling Technology | Cat.# 74512 |
| Anti-USP50 | Affinity | Cat.# AF9225<br>RRID: AB_2843415 |
| Anti-USP50 | Proteintech | Cat.# 20374-1-AP<br>RRID: AB_10973680 |
| Anti-Myc | Abmart | Cat.# M20002H |
| Anti-Flag | Sigma | Cat.# F7425<br>RRID: AB_439687 |
| Anti-HA | Abcam | Cat.# ab9110<br>RRID: AB_307019 |
| Anti-STAT1 | Cell Signaling Technology | Cat.# 9172<br>RRID: AB_2198300 |
| Anti-Tyk2 | Cell Signaling Technology | Cat.# 14193<br>RRID: AB_2798419 |
| Anti-VSV-G | Santa Cruz | Cat.# sc-66180<br>RRID: AB_832793 |
| Anti-β-Actin | Proteintech | Cat.# 66009-1-Ig<br>RRID: AB_2687938 |
| Anti-Tubulin | Proteintech | Cat.# 66031-1-Ig<br>RRID: AB_11042766 |
| Anti-Ubiquitin (Ub) | Santa Cruz | Cat.# sc-8017<br>RRID: AB_2762364 |
| Anti-K48 Ub | Cell Signaling Technology | Cat.# 4289S<br>RRID: AB_10557239 |
| Anti-IRF3 | Santa Cruz | Cat.# sc-33641<br>RRID: AB_627826 |
| Anti-SARS-CoV-2-Spike | GeneTex | Cat.# GTX632604<br>RRID: AB_2864418 |
| Bacterial and Virus Strains |  |  |
| DH5α | Transgen Biotech | Cat # CD201 |
| SARS-CoV-2 GFP/ΔN | Dr. Qiang Ding from<br>Tsinghua University, China | N/A |
| SARS-CoV-2-S | Dr. Aihua Zheng from<br>Chinese Academy of<br>Sciences | N/A |
| VSV | Dr. Chen Wang from China<br>Pharmaceutical University | N/A |
| VSV-GFP | Dr. Chunsheng Dong from<br>Soochow University, China | N/A |
| Chemicals, and Recombinant Proteins |  |  |
| Flag peptide | Sigma | Cat.# F3290 |
| TRIzol reagent | TAKARA | Cat.# 9109 |
| Vitamin C (VitC) | Sigma | Cat.# A7506 |
| r-hACE2-IgG-Fc | GenScript | Z03516 |
| Critical Commercial Assays |  |  |
| QuickChange Lightning<br>site-Directed Mutagenesis<br>Kit | TIANGEN | Cat.# KM101 |
| Vitamin-C Detection Kit | Elabscience | E-BC-K034-S |
| Experimental Models: Cell Lines |  |  |
| HEK293T | American Type Culture<br>Collection | N/A |
| A549 | American Type Culture<br>Collection | N/A |
| 2fTGH | Dr. Serge Y. Fuchs from | N/A |
|  | University of Pennsylvania,<br>America |  |
| Caco-2-N | Dr. Qiang Ding from<br>Tsinghua University, China | N/A |
| Experimental Models: Organisms/Strains |  |  |
| hACE2 mice | Cyagen Biosciences | N/A |
| C57BL/6 mice | Shanghai SLAC<br>Laboratory Animals | N/A |

## CONTACT FOR REAGENT AND RESOURCE SHARING

Further information and requests for resources and reagents should be directed to and will be fulfilled by the Lead Contact, Hui Zheng.

## EXPERIMENTAL MODEL AND SUBJECT DETAILS

### Mice

Mice with human *Ace2* gene in C57BL/6 background (hACE2 mice) were generated by the Cyagen Biosciences *Inc*. (Guangzhou). Wild-type (WT) C57BL/6 mice were purchased from the Shanghai SLAC Laboratory Animals. All mice were maintained under specific-pathogen-free (SPF) conditions in the animal facility of Soochow University. We used 6-8 weeks old mice in all experiments. Animal care and use protocol adhered to the National Regulations for the Administration of Affairs Concerning Experimental Animals. All animal experiments have received ethical approval by the Ethics Committee of the Soochow University, and were carried out in accordance with the Laboratory Animal Management Regulations with approval of the Scientific Investigation Board of Soochow University, Suzhou.

## METHOD DETAILS

### Cell culture and reagents

HEK293T, Caco-2 and A549 cells were obtained from ATCC. 2fTGH cells were gifts from Dr. Serge Y. Fuchs (University of Pennsylvania). Caco-2-N cells that stably express a SARS-CoV-2 nucleocapsid (N) gene were gifts from Dr. Qiang Ding (Tsinghua University). All cells were cultured at 37 °C under 5% CO_2_ in DMEM (HyClone) supplemented with 10% FBS (GIBCO, Life Technologies), 100 unit/ml penicillin, and 100 µg/ml streptomycin. Vitamin B1, B6, B12, C, D3 and K1 were from Sigma. VitC-Na^+^ was made by Dr. Yibo Zuo (Soochow University). Flag peptides (F3290), puromycin and other chemicals were purchased from Sigma.

### Plasmids and transfection

Flag-ACE2 plasmid was purchased from the Beyotime Biotechnology (D2949). Myc-ACE2 and its deletion mutants were generated using PCR amplified from Flag-ACE2 or Myc-ACE2. Flag-HA (FH)-tagged human DUBs, including FH-USP50, were gifts from Dr. J. Wade Harper (Harvard Medical School, Addgene plasmids). HA-Ub (K6, K11, K27, K29, K33, K48 and K63) were gifts from Dr. Lingqiang Zhang (State Key Laboratory of Proteomics, China). All plasmids were confirmed by sequencing. The shACE2 and shUSP50 was constructed into the shX vector that is a gift from Dr. Jianfeng Dai (Soochow University) with following sequences:

shACE2 (#1): GGACAAGTTTAACCACGAAGC;

shACE2 (#2): GCAAACGGTTGAACACAATTC;

shUSP50 (#1): CCCGGAGAAGATCATATGA;

shUSP50 (#2): GTTTGAAGAGCAGCTCAAT.

All mutations were generated by the QuickChange Lightning site-Directed Mutagenesis Kit (TIANGEN, KM101). Transient transfections for different cell lines were carried out using the LongTrans (Ucallm, TF/07).

### Liver, lung and kidney histology

Liver, lung and kidney tissues from mice administrated with or without VitC (300 mg/kg of body weight) were fixed in 4% formaldehyde solution, and then embedded into paraffin. Briefly, the paraffin sections were stained with anti-human ACE2 antibodies, and then observed by light microscopy for histological changes. In addition, mice were administrated with VitC (300 mg/day/kg of body weight) for two days and then infected with SARS-CoV-2-S pseudoviruses (1×10^6^ PFU per gram body) for 24 hrs. Mouse lung and kidney tissues were stained using antibodies against SARS-CoV-2-Spike (Genetex, GTX632604). DAPI was used for nuclei staining. Representative images are shown at 100× magnification.

### VitC detection assay

HEK293T cells were transfected with either Flag-USP50 or Myc-ACE2 (WT or its deletion mutants). After 48 hrs, Flag-USP50 or Myc-ACE2 proteins were immunoprecipitated by either Flag (M2) beads or a Myc antibody with Protein-G beads at 4 °C. After washing three times, the immunoprecipitates (Flag-USP50 proteins with Flag beads or Myc-ACE2 proteins with Protein-G beads) were mixed with VitC for binding for 2 hrs at 4°C. After centrifuging, the supernatant was discarded, and the precipitates were washing and then diluted in 1× PBS buffer for further determination of VitC concentrations by the Vitamin-C Detection Kit (Elabscience, E-BC-K034-S).

In addition, recombinant human ACE2-IgG-Fc fragments (r-hACE2-Fc) (GenScript, Z03516) were firstly incubated with the Protein-G agarose (Millipore, 16-266) for 2 hrs. An anti-Tyk2 IgG protein (Cell Signaling Technology, 14193) was used as a control for the VitC binding experiment. After washing and centrifuging, the pellets were diluted in 1× PBS buffer and then mixed with VitC for 2 hrs at 4 °C for binding. VitC concentrations were measured by the Vitamin-C Detection Kit (Elabscience, E-BC-K034-S).

### *In vitro* protein-protein binding assay

HEK293T cells were transfected with either Flag-USP50 or Myc-ACE2. Flag-USP50 proteins were immunoprecipitated by Flag (M2) beads and then eluted with the Flag peptides. Myc-ACE2 was pulled down by the Myc antibody from HEK293T cells transfected with Myc-ACE2. Next, Flag-USP50 eluates and the immunoprecipitation beads with Myc-ACE2 proteins were mixed with or without VitC, and then vibrated for 2 hrs at 4 °C. After washing and centrifuging, SDS-PAGE and immunoblotting were used to analyze Flag-USP50 and Myc-ACE2 protein levels using anti-Flag or anti-Myc antibodies.

### CRISPR-Cas9 genome editing

The lenti-CRISPRv2 vector was a nice gift from Dr. Fangfang Zhou (Soochow University, China). For gene knockout, small guide RNAs were firstly cloned into the lenti-CRISPRv2 vector, and then transfected into HEK293T cells. Forty-eight hours after transfection, the cells were cultured under puromycin (1.5 µg/ml) selection for 2 weeks, and then cells were identified by immunoblotting analysis. After that, cells were transferred to 96-well plates and cultured for further experiments. The guide RNA sequences are as following: human *Usp50*: 5′-CCCCATTTTCAGGGTGTCAC-3′.

### VitC docking with ACE2

VitC docking with ACE2 was performed by Prof. Sheng Tian (Soochow University). The binding pocket is defined by the centroid of amino acid residues 200-400 of ACE2. Based on the glide docking module in the Schrödinger molecular simulation software package, VitC was aligned to the defined binding pocket using standard precision (SP), and the binding conformation and its interaction were evaluated. The binding conformation and interaction diagram of VitC and ACE2 are obtained based on SP docking scoring function.

### Immunoblotting (IB) and Immunoprecipitation (IP)

Cells were harvested using the lysis buffer containing 1% Nonidet P-40 (NP-40), 150 mM NaCl, Tris-HCl (20 mM, pH 7.4), 0.5 mM EDTA, PMSF (50 µg/ml) and protease inhibitor mixtures (Sigma). Proteins from whole cell lysates were firstly subjected to SDS-PAGE, and then transferred to PVDF membranes (Millipore). After blocking with 5% nonfat milk for 1 hr, the membranes were incubated with the corresponding primary antibodies overnight, followed by incubation with the secondary antibodies (Bioworld or Abbkine). All immunoreactive bands were visualized with the NcmECL Ultra (MCM Biotech, P10300).

Immunoprecipitation was firstly carried out using specific antibodies at 4 °C. Protein G agarose beads (Millipore, #16-266) were then added and incubated for 2 hrs on a rotor at 4°C. After washing five times with the lysis buffer, the immunoprecipitates were eluted by heating at 95 °C with the loading buffer containing β-mercaptoethanol for 10 min and then analyzed by SDS-PAGE gels and subsequent immunoblotting.

The antibodies with the indicated dilutions were as follows: anti-human ACE2 (Cell Signaling Technology, #74512, 1:1000), anti-ACE2 (Affinity, AF5165, 1:1000), anti-USP50 (Affinity, AF9225, 1:1000), anti-USP50 (Proteintech, 20374-1-AP, 1:1000), anti-Flag (Sigma, F7425, 1:5000), anti-HA (Abcam, ab9110, 1:3000), anti-Myc (Abmart, M20002H, 1:3000), anti-IRF3 (Santa Cruz, sc-33641, 1:1000), anti-STAT1 (Cell Signaling Technology, 9172, 1:1000), anti-Ubiquitin (Ub) (Santa Cruz, 12987-1-AP, 1:1000), anti-K48 Ub (Cell Signaling Technology, 4289S, 1:1000), anti-VSV-G (Santa Cruz, sc-66180, 1:2000) and anti-Tubulin (Proteintech, 66031-1-Ig, 1:3000).

### RNA isolation and quantitative real-time PCR

Total RNAs were isolated from different cells or mouse tissues using a TRIzol reagent (Invitrogen). The detailed procedures for RNA isolation and quantitative real-time PCR (RT-qPCR) were as described previously (Zuo et al., 2020; Zuo et al., 2022). Briefly, the cDNA was synthesized using 5× All-In-One RT MasterMix (abm, #G490, Beijing). RT-qPCR was performed using a StepOne Plus real-time PCR system (Applied Bioscience). The results from three independent experiments were shown as the average mean ± standard deviation (s.d.). The primer sequences are as following:

human *Ace2*:

Forward: 5’-ACCACGAAGCCGAAGACCTGTT-3’

Reverse: 5’-TGGGCAAGTGTGGACTGTTCCT-3’;

SARS-CoV-2 GFP/ΔN:

Forward: 5’-GCTTTGCTGGAAATGCCGTT-3’

Reverse: 5’-GGACTTGTTGTGCCATCACC-3’;

SARS-CoV-2-S:

Forward: 5’-ATGTCCTTCCCTCAGTCAGCAC-3’

Reverse: 5’-TGACAAATGGCAGGAGCAGTTG-3’; VSV:

Forward: 5’-ACGGCGTACTTCCAGATGG-3’

Reverse: 5’-CTCGGTTCAAGATCCAGGT-3’;

β*-actin*:

Forward: 5’-ACCAACTGGGACGACATGGAGAAA-3’

Reverse: 5’-ATAGCACAGCCTGGATAGCAACG-3’.

### Virus and viral infection *in vitro*

SARS-CoV-2 GFP/ΔN viruses were gifts from Dr. Qiang Ding (Tsinghua University) (Ju et al., 2021). SARS-CoV-2-S pseudoviruses, which contain the rVSV-eGFP-SARS-CoV-2 backbone with the VSV glycoprotein coding sequence (3845–5380) being replaced by the SARS-CoV-2 Spike gene (Li et al., 2020), were nice gifts from Dr. Aihua Zheng (Chinese Academy of Sciences) and Dr. Jianfeng Dai (Soochow University, China). Vesicular stomatitis virus (VSV) was a gift from Dr. Chen Wang (China Pharmaceutical University). VSV-GFP viruses were gifts from Dr. Chunsheng Dong (Soochow University, China). Cells were treated with VitC (5 mM) overnight. After washing twice, cells were infected by either SARS-CoV-2, or VSV, or VSV-GFP at a multiplicity of infection (MOI) of 0.1 for 24 hrs. Then cells were analyzed by immunofluorescence, RT-qPCR or western blot.

### Viral infection *in vivo*

For *in vivo* viral infection studies, 8-week-old hACE2 mice were intraperitoneally administrated with Vitamin C (300 mg/kg body weight, once a day) for two days. Then mice were given intraperitoneal injections (*i*.*p*.) of SARS-CoV-2-S pseudoviruses (1×10^6^ PFU per gram body mouse). Twenty-four hours after infection, mouse lung, liver, kidney and spleen tissues were harvested. RT-qPCR was performed for the analysis of SARS-CoV-2-S viral RNA levels.

### Immunofluorescence microscopy

HeLa cells were firstly washed by 1x PBS and fixed in 4% paraformaldehyde on ice. Then cells were permeabilized with Triton X-100 (0.5%) and blocked with BSA (5%). Next, cells were incubated with two different anti-ACE2 antibodies overnight in 0.5% BSA. After washing three times with 1x PBS, cells were stained with either 488 goat anti-mouse IgG (Alexa Fluor, A11001) or 594 goat anti-rabbit IgG (Alexa Fluor, A11012). Nucleus were stained with DAPI. The fluorescent images were captured with the Nikon A1 confocal microscope.

### QUANTIFICATION STATISTICAL ANALYSIS

#### Statistical analysis

Two-tailed unpaired Student’s *t*-test was employed to compare the significance between different groups. All differences were considered statistically significant when *p* < 0.05. P values are indicated by asterisks in the figures as follows: \**p* < 0.05, \*\**p* < 0.01 and \*\*\**p* < 0.001.

## DATA AND SOFTWARE AVAILABILITY

All data generated or analyzed during this study are included in the Figures 1-6 and Supplementary Figures 1-7. Additional datasets that support the findings of this study are available from the corresponding author upon reasonable request.

## SUPPLEMENTAL INFORMATION

Supplemental Information includes seven figures and can be found with this article online.

## ACKNOWLEDGMENTS

We thank Dr. Serge Y. Fuchs (University of Pennsylvania), Dr. J. Wade Harper (Harvard Medical School), Dr. Lingqiang Zhang (National Center of Protein Sciences, China), Dr. Chen Wang (China Pharmaceutical University), Dr. Chunsheng Dong (Soochow University, China) and Dr. Aihua Zheng (Chinese Academy of Sciences) for important reagents. This work is supported by the National Natural Science Foundation of China (32100568, 31970846), the National Key R&D Program of China (2018YFC1705500): 2018YFC1705505, the Postdoctoral Science Foundation of China (2021M692351), and the Priority Academic Program Development of Jiangsu Higher Education Institutions (PAPD).

## AUTHOR CONTRIBUTIONS

Y.Z., Z.Z., Y.H., J.H., L.Z., T.R., X.C and Y.M. performed the experiments. Z.Z., Y.H. and F.M. assisted with mouse experiments, tissue processing and analysis. S.T. performed VitC docking analysis. Q.D. assisted with SARS-CoV-2 GFP/ΔN experiments. H.Z. and Y.Z. designed experiments, analyzed data and wrote the paper. Y.Y., Y.L., J.D., Q.D., S.T. and H.Z. discussed the data and manuscript. H.Z. was responsible for research supervision, coordination, and strategy.

## REFERENCES

Ju, X., Zhu, Y., Wang, Y., Li, J., Zhang, J., Gong, M., Ren, W., Li, S., Zhong, J., Zhang, L., et al. (2021). A novel cell culture system modeling the SARS-CoV-2 life cycle. PLoS Pathog 17, e1009439.

Li, H., Zhao, C., Zhang, Y., Yuan, F., Zhang, Q., Shi, X., Zhang, L., Qin, C., and Zheng, A. (2020). Establishment of replication-competent vesicular stomatitis virus-based recombinant viruses suitable for SARS-CoV-2 entry and neutralization assays. Emerg Microbes Infect 9, 2269–2277.

Zuo, Y., Feng, Q., Jin, L., Huang, F., Miao, Y., Liu, J., Xu, Y., Chen, X., Zhang, H., Guo, T., et al. (2020). Regulation of the linear ubiquitination of STAT1 controls antiviral interferon signaling. Nat Commun 11, 1146.

Zuo, Y., He, J., Liu, S., Xu, Y., Liu, J., Qiao, C., Zang, L., Sun, W., Yuan, Y., Zhang, H., et al. (2022). LATS1 is a central signal transmitter for achieving full type-I interferon activity. Sci Adv 8, eabj3887.

## REFERENCES

Cai, J., Wei, J., Schrott, V., Zhao, J., Bullock, G., and Zhao, Y. (2018). Induction of deubiquitinating enzyme USP50 during erythropoiesis and its potential role in the regulation of Ku70 stability. J Investig Med 66, 1–6.

Chavda, V.P., Prajapati, R., Lathigara, D., Nagar, B., Kukadiya, J., Redwan, E.M., Uversky, V.N., Kher, M.N., and Patel, R. (2022). Therapeutic monoclonal antibodies for COVID-19 management: an update. Expert Opin Biol Ther 22, 763–780.

Chen, Q., Espey, M.G., Krishna, M.C., Mitchell, J.B., Corpe, C.P., Buettner, G.R., Shacter, E., and Levine, M. (2005). Pharmacologic ascorbic acid concentrations selectively kill cancer cells: action as a pro-drug to deliver hydrogen peroxide to tissues. Proc Natl Acad Sci U S A 102, 13604–13609.

Crackower, M.A., Sarao, R., Oudit, G.Y., Yagil, C., Kozieradzki, I., Scanga, S.E., Oliveira-dos-Santos, A.J., da Costa, J., Zhang, L., Pei, Y., et al. (2002). Angiotensin-converting enzyme 2 is an essential regulator of heart function. Nature 417, 822–828.

Donoghue, M., Hsieh, F., Baronas, E., Godbout, K., Gosselin, M., Stagliano, N., Donovan, M., Woolf, B., Robison, K., Jeyaseelan, R., et al. (2000). A novel angiotensin-converting enzyme-related carboxypeptidase (ACE2) converts angiotensin I to angiotensin 1-9. Circ Res 87, E1–9.

Edenfield, R.C., and Easley, C.A.t. (2022). Implications of testicular ACE2 and the renin-angiotensin system for SARS-CoV-2 on testis function. Nat Rev Urol 19, 116–127.

Ferrario, C.M., Jessup, J., Chappell, M.C., Averill, D.B., Brosnihan, K.B., Tallant, E.A., Diz, D.I., and Gallagher, P.E. (2005). Effect of angiotensin-converting enzyme inhibition and angiotensin II receptor blockers on cardiac angiotensin-converting enzyme 2. Circulation 111, 2605–2610.

Garvin, M.R., Alvarez, C., Miller, J.I., Prates, E.T., Walker, A.M., Amos, B.K., Mast, A.E., Justice, A., Aronow, B., and Jacobson, D. (2020). A mechanistic model and therapeutic interventions for COVID-19 involving a RAS-mediated bradykinin storm. Elife 9.

Guan, W.J., Ni, Z.Y., Hu, Y., Liang, W.H., Ou, C.Q., He, J.X., Liu, L., Shan, H., Lei, C.L., Hui, D.S.C., et al. (2020). Clinical Characteristics of Coronavirus Disease 2019 in China. N Engl J Med 382, 1708–1720.

Gupta, A., Madhavan, M.V., Sehgal, K., Nair, N., Mahajan, S., Sehrawat, T.S., Bikdeli, B., Ahluwalia, N., Ausiello, J.C., Wan, E.Y., et al. (2020). Extrapulmonary manifestations of COVID-19. Nat Med 26, 1017–1032.

Hoffmann, M., Kleine-Weber, H., Schroeder, S., Kruger, N., Herrler, T., Erichsen, S., Schiergens, T.S., Herrler, G., Wu, N.H., Nitsche, A., et al. (2020). SARS-CoV-2 Cell Entry Depends on ACE2 and TMPRSS2 and Is Blocked by a Clinically Proven Protease Inhibitor. Cell 181, 271–280 e278.

Krege, J.H., John, S.W., Langenbach, L.L., Hodgin, J.B., Hagaman, J.R., Bachman, E.S., Jennette, J.C., O’Brien, D.A., and Smithies, O. (1995). Male-female differences in fertility and blood pressure in ACE-deficient mice. Nature 375, 146–148.

Kuba, K., Imai, Y., and Penninger, J.M. (2006). Angiotensin-converting enzyme 2 in lung diseases. Curr Opin Pharmacol 6, 271–276.

Levine, M., Conry-Cantilena, C., Wang, Y., Welch, R.W., Washko, P.W., Dhariwal, K.R., Park, J.B., Lazarev, A., Graumlich, J.F., King, J., et al. (1996). Vitamin C pharmacokinetics in healthy volunteers: evidence for a recommended dietary allowance. Proc Natl Acad Sci U S A 93, 3704–3709.

Li, M.Y., Li, L., Zhang, Y., and Wang, X.S. (2020). Expression of the SARS-CoV-2 cell receptor gene ACE2 in a wide variety of human tissues. Infect Dis Poverty 9, 45.

Li, R., Li, X., Zhao, J., Meng, F., Yao, C., Bao, E., Sun, N., Chen, X., Cheng, W., Hua, H., et al. (2022). Mitochondrial STAT3 exacerbates LPS-induced sepsis by driving CPT1a-mediated fatty acid oxidation. Theranostics 12, 976–998.

Lindblad, M., Tveden-Nyborg, P., and Lykkesfeldt, J. (2013). Regulation of vitamin C homeostasis during deficiency. Nutrients 5, 2860–2879.

Lykkesfeldt, J., and Tveden-Nyborg, P. (2019). The Pharmacokinetics of Vitamin C. Nutrients 11.

Messerli, F.H., Bangalore, S., Bavishi, C., and Rimoldi, S.F. (2018). Angiotensin-Converting Enzyme Inhibitors in Hypertension: To Use or Not to Use? J Am Coll Cardiol 71, 1474–1482.

Mohamed, T., Abdul-Hafez, A., and Uhal, B.D. (2021). Regulation of ACE-2 enzyme by hyperoxia in lung epithelial cells by post-translational modification. J Lung Pulm Respir Res 8, 47–52.

Nielsen, T.K., Hojgaard, M., Andersen, J.T., Poulsen, H.E., Lykkesfeldt, J., and Mikines, K.J. (2015). Elimination of ascorbic acid after high-dose infusion in prostate cancer patients: a pharmacokinetic evaluation. Basic Clin Pharmacol Toxicol 116, 343–348.

Padayatty, S.J., Sun, H., Wang, Y., Riordan, H.D., Hewitt, S.M., Katz, A., Wesley, R.A., and Levine, M. (2004). Vitamin C pharmacokinetics: implications for oral and intravenous use. Ann Intern Med 140, 533–537.

Ritorto, M.S., Ewan, R., Perez-Oliva, A.B., Knebel, A., Buhrlage, S.J., Wightman, M., Kelly, S.M., Wood, N.T., Virdee, S., Gray, N.S., et al. (2014). Screening of DUB activity and specificity by MALDI-TOF mass spectrometry. Nat Commun 5, 4763.

Rodrigues Prestes, T.R., Rocha, N.P., Miranda, A.S., Teixeira, A.L., and Simoes, E.S.A.C. (2017). The Anti-Inflammatory Potential of ACE2/Angiotensin-(1-7)/Mas Receptor Axis: Evidence from Basic and Clinical Research. Curr Drug Targets 18, 1301–1313.

Rothlin, R.P., Vetulli, H.M., Duarte, M., and Pelorosso, F.G. (2020). Telmisartan as tentative angiotensin receptor blocker therapeutic for COVID-19. Drug Dev Res 81, 768–770.

Shen, H., Zhang, J., Wang, C., Jain, P.P., Xiong, M., Shi, X., Lei, Y., Chen, S., Yin, Q., Thistlethwaite, P.A., et al. (2020). MDM2-Mediated Ubiquitination of Angiotensin-Converting Enzyme 2 Contributes to the Development of Pulmonary Arterial Hypertension. Circulation 142, 1190–1204.

Stephenson, C.M., Levin, R.D., Spector, T., and Lis, C.G. (2013). Phase I clinical trial to evaluate the safety, tolerability, and pharmacokinetics of high-dose intravenous ascorbic acid in patients with advanced cancer. Cancer Chemother Pharmacol 72, 139–146.

Uhler, C., and Shivashankar, G.V. (2020). Mechano-genomic regulation of coronaviruses and its interplay with ageing. Nat Rev Mol Cell Biol 21, 247–248.

Verdecchia, P., Angeli, F., Mazzotta, G., Ambrosio, G., and Reboldi, G. (2010). Angiotensin converting enzyme inhibitors and angiotensin receptor blockers in the treatment of hypertension: should they be used together? Curr Vasc Pharmacol 8, 742–746.

Vilcheze, C., Kim, J., and Jacobs, W.R., Jr. (2018). Vitamin C Potentiates the Killing of Mycobacterium tuberculosis by the First-Line Tuberculosis Drugs Isoniazid and Rifampin in Mice. Antimicrob Agents Chemother 62.

Wang, G., Zhao, Q., Zhang, H., Liang, F., Zhang, C., Wang, J., Chen, Z., Wu, R., Yu, H., Sun, B., et al. (2021). Degradation of SARS-CoV-2 receptor ACE2 by the E3 ubiquitin ligase Skp2 in lung epithelial cells. Front Med 15, 252–263.

Wang, L., Wu, Y., Yao, S., Ge, H., Zhu, Y., Chen, K., Chen, W.Z., Zhang, Y., Zhu, W., Wang, H.Y., et al. (2022). Discovery of potential small molecular SARS-CoV-2 entry blockers targeting the spike protein. Acta Pharmacol Sin 43, 788–796.

Winkelmayer, W.C., Fischer, M.A., Schneeweiss, S., Wang, P.S., Levin, R., and Avorn, J. (2005). Underuse of ACE inhibitors and angiotensin II receptor blockers in elderly patients with diabetes. Am J Kidney Dis 46, 1080–1087.

Wrapp, D., Wang, N., Corbett, K.S., Goldsmith, J.A., Hsieh, C.L., Abiona, O., Graham, B.S., and McLellan, J.S. (2020). Cryo-EM structure of the 2019-nCoV spike in the prefusion conformation. Science 367, 1260–1263.

Xu, Y., Wu, C., Cao, X., Gu, C., Liu, H., Jiang, M., Wang, X., Yuan, Q., Wu, K., Liu, J., et al. (2022). Structural and biochemical mechanism for increased infectivity and immune evasion of Omicron BA.2 variant compared to BA.1 and their possible mouse origins. Cell Res.

Ziegler, C.G.K., Allon, S.J., Nyquist, S.K., Mbano, I.M., Miao, V.N., Tzouanas, C.N., Cao, Y., Yousif, A.S., Bals, J., Hauser, B.M., et al. (2020). SARS-CoV-2 Receptor ACE2 Is an Interferon-Stimulated Gene in Human Airway Epithelial Cells and Is Detected in Specific Cell Subsets across Tissues. Cell 181, 1016–1035 e1019.

