## Supplementary Figures 1-7 for "Vitamin C is an efficient natural product for prevention of SARS-CoV-2 infection by targeting ACE2 in both cell and in vivo mouse models"

**This PDF file includes:**

Figs. S1 to S7

### Supplementary Figure 1

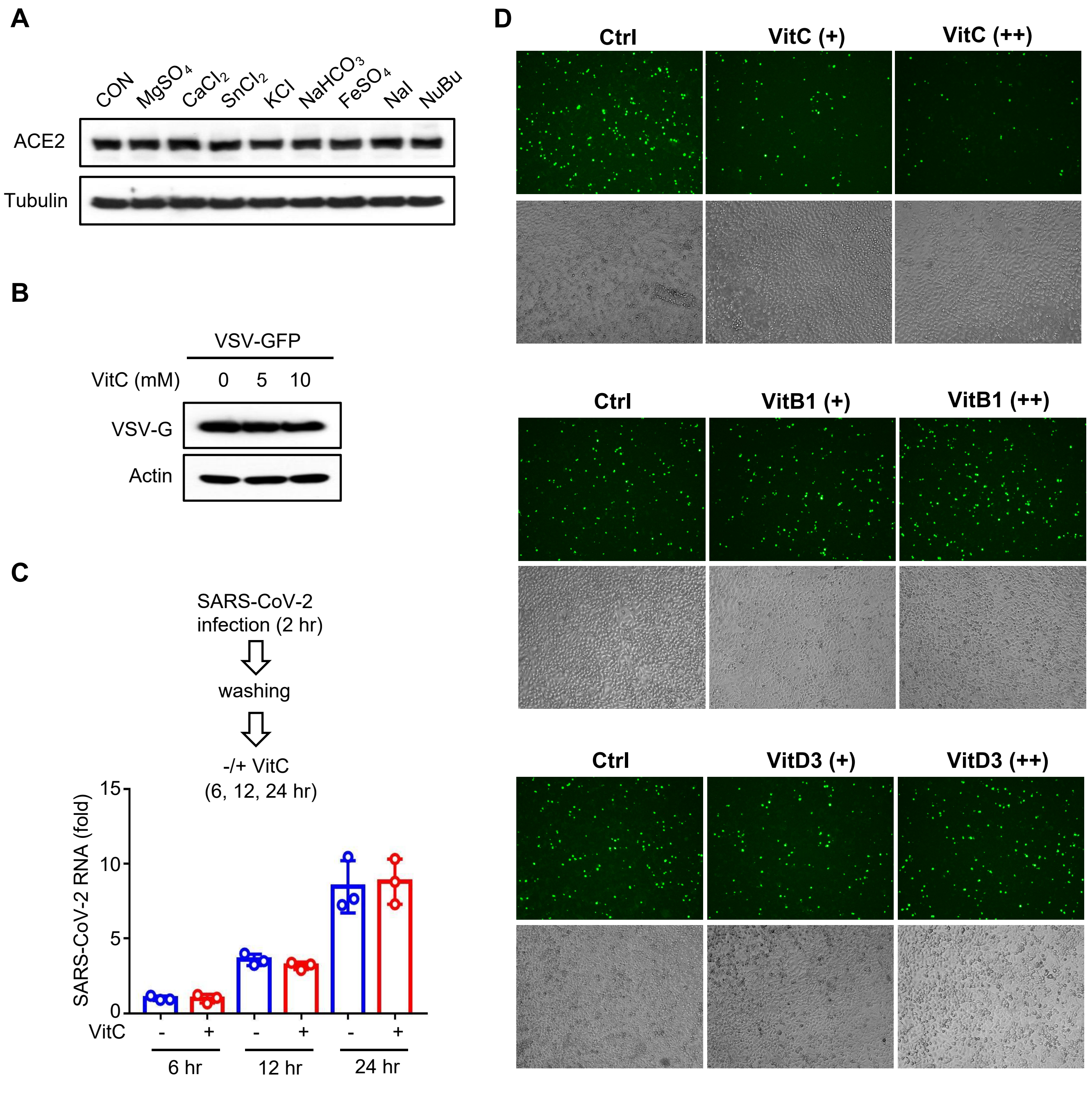

**Figure S1. VitC restricts cellular infection with the SARS-CoV-2-S pseudovirus, Related to Figure 1**

**(A)** Western blot analysis of ACE2 in Caco-2 cells treated with 20 μM of individual compounds (MgSO4, CaCl2, SnCl2, KCl, NaHCO3, FeSO4, NaI and NuBu) for 24 hrs.

**(B)** Western blot analysis of VSV-G proteins in Caco-2 cells pretreated with Vitamin C (VitC, 5 mM) for 24 hrs and then infected with VSV-GFP (MOI = 0.1) for 24 hrs.

**(C)** Caco-2-N cells were infected with SARS-CoV-2 GFP/ΔN (MOI = 0.1) for 2 hrs. After washing, cells were treated with VitC (2.5 mM) for 6, 12 and 24 hrs. RT-qPCR was used to analyze SARS-CoV-2 RNA levels.

**(D)** Fluorescence microscopy analysis of the SARS-CoV-2-S pseudovirus in HEK293T cells pretreated with VitC (2.5 mM and 5 mM) or Vitamin B1 (VitB1, 250 μM and 500 μM) or Vitamin D3 (VitD3, 12.5 μM and 25 μM) for 12 hrs, followed by infection with the SARS-CoV-2-S GFP pseudovirus (MOI = 0.1) for 24 hrs.

Data are representative of three independent experiments (A, B), or are shown as mean and s.d. of three biological replicates (C). N.S, not significant ($p > 0.05$), two-tailed unpaired Student's *t*-test.

### Supplementary Figure 2

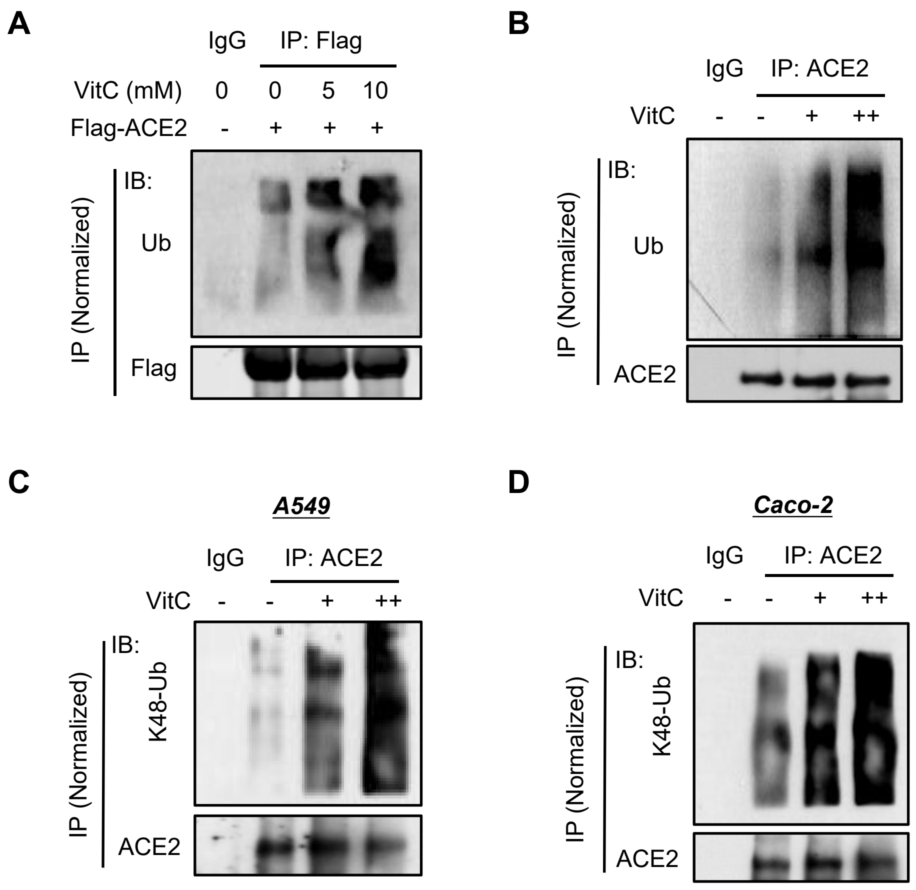

**Figure S2. VitC treatment upregulates K48-linked polyubiquitination of ACE2, Related to Figure 2**

**(A)** Immunoprecipitation (IP)-immunoblotting (IB) analysis of total ubiquitination of Flag-ACE2 in HEK293T cells transfected with Flag-ACE2 and then treated with VitC as indicated for 12 hrs.

**(B)** IP-IB analysis of total ubiquitination of endogenous ACE2 in Caco-2 cells treated with VitC (2.5 mM and 5 mM) for 12 hrs.

**(C, D)** IP-IB analysis of K48-linked polyubiquitination of endogenous ACE2 in A549 (C) and Caco-2 (D) cells treated with VitC (2.5 mM and 5 mM) for 12 hrs.

Data are representative of three independent experiments (A-D).

### Supplementary Figure 3

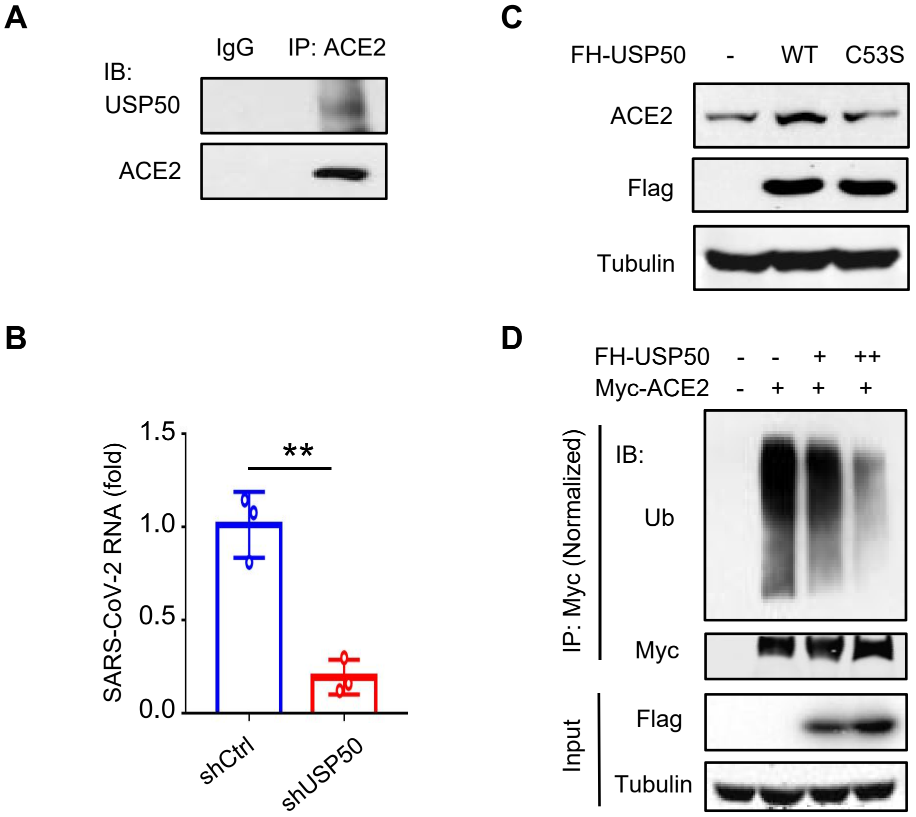

**Figure S3. USP50 interacts with ACE2 and regulates ACE2 protein levels dependently on its deubiquitinase activity, Related to Figure 3**

**(A)** IP-IB analysis of the interaction between endogenous USP50 and ACE2 in HEK293T cells.

**(B)** RT-qPCR analysis of SARS-CoV-2 GFP/ΔN RNA levels in 2fTGH cells transfected with control shRNAs (shCtrl) or shRNAs against USP50 (shUSP50) and then infected with SARS-CoV-2 GFP/ΔN (MOI = 0.1) for 24 hrs.

**(C)** Western blot analysis of ACE2 in HEK293T cells transfected with Flag-HA-tagged USP50 (FH-USP50) wild-type (WT) or its deubiquitinase inactive mutant (C53S).

**(D)** IP-IB analysis of ubiquitination of Myc-ACE2 in HEK293T cells cotransfected with Myc-ACE2 and increasing amounts of Flag-USP50.

Data are representative of three independent experiments (A, C, D), or are shown as mean and s.d. of three biological replicates (B). $^{**}p < 0.01$ (two-tailed unpaired Student's *t*-test).

### Supplementary Figure 4

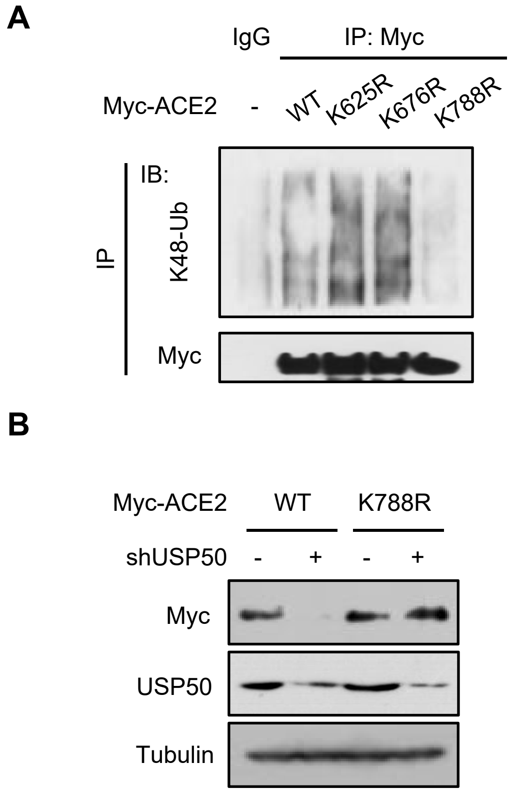


**Figure S4. USP50 regulates ACE2 dependently on the Lys788 residue of ACE2, Related to Figure 4**

**(A)** IP-IB analysis of K48-linked polyubiquitination of Myc-ACE2 in HEK293T cells transfected with Myc-ACE2 (WT or its mutants).

**(B)** Western blot analysis of Myc-ACE2 in HEK293T cells transfected with Myc-ACE2 (WT or K788R), together with shCtrl (-) or shUSP50 (+).

Data are representative of three independent experiments (A, B).

### Supplementary Figure 5

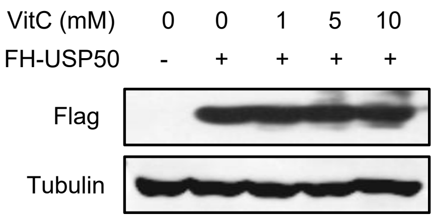


**Figure S5. VitC does not substantially affect USP50 protein levels, Related to Figure 5**

Western blot analysis of FH-USP50 levels in HEK293T cells transfected with FH-USP50 and then treated with VitC as indicated for 12 hrs.

Data are representative of three independent experiments.

### Supplementary Figure 6

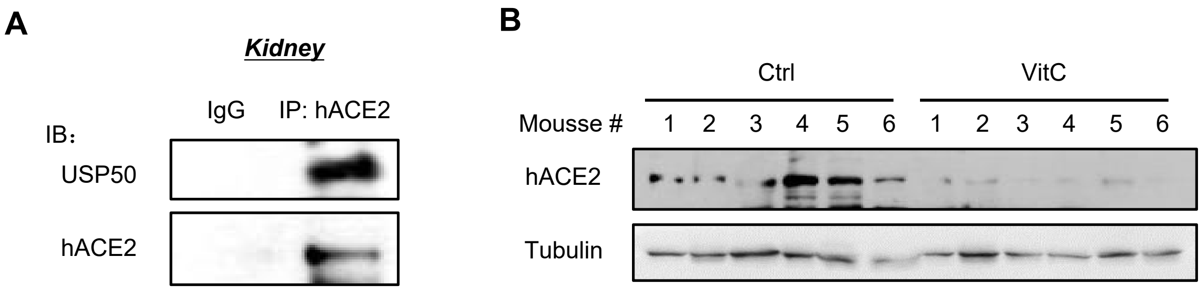


**Figure S6. USP50 interacts with hACE2 *in vivo*, Related to Figure 6**

**(A)** IP-IB analysis of the interaction between endogenous hACE2 and USP50 in kidney tissues of hACE2 mice.

**(B)** The hACE2 mice were intraperitoneally administrated with VitC (300 mg/day/kg body weight) for two days. Western blot analysis of hACE2 levels in spleen tissues of hACE2 mice.

Data are representative of three independent experiments (A, B).

### Supplementary Figure 7

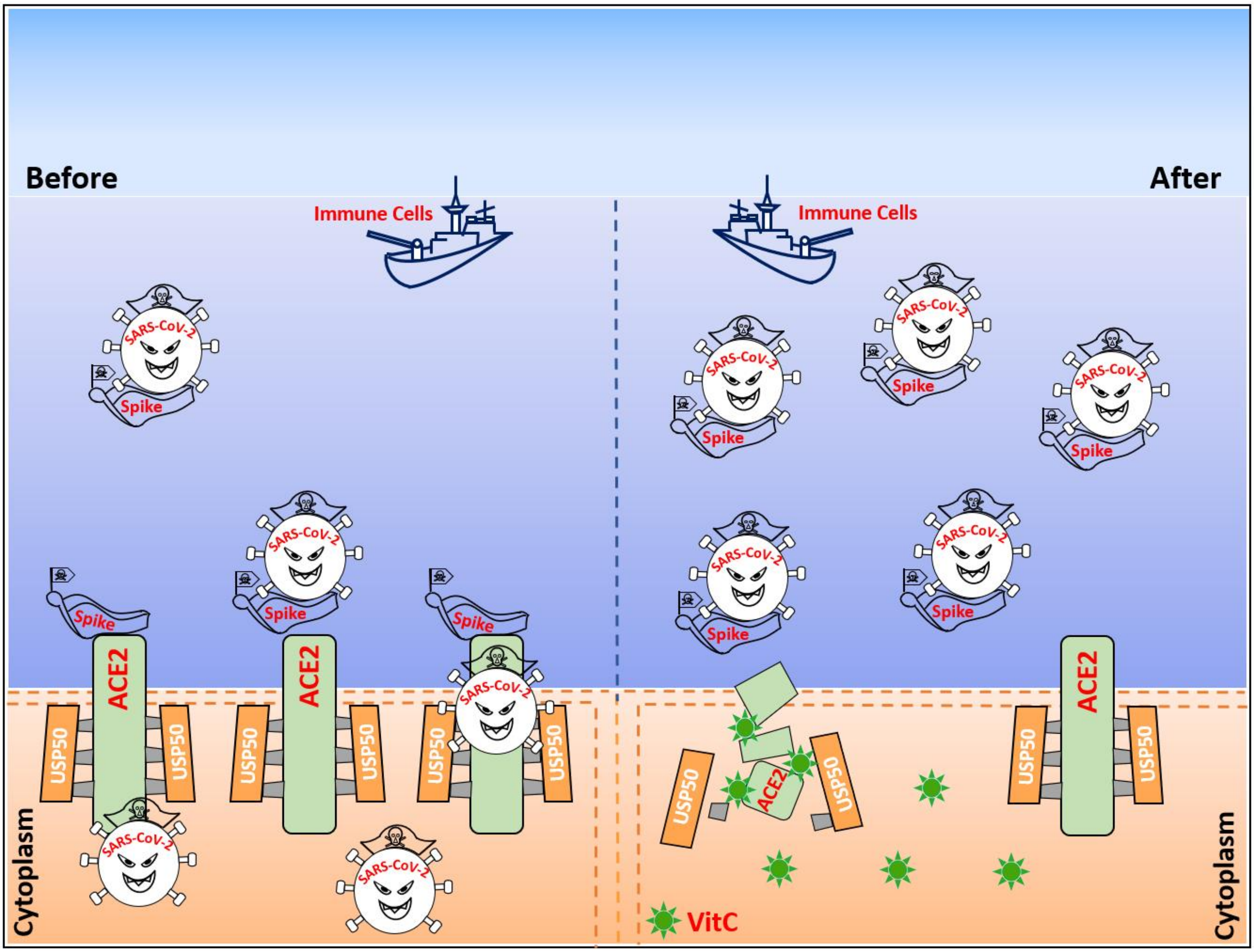


**Figure S7. A proposed model of VitC-regulated SARS-CoV-2 infection**

The deubiquitinase USP50 controls ACE2 protein stability and levels, while Vitamin C blocks the USP50-ACE2 interaction and therefore results in ACE2 degradation, offering a flexible and efficient approach to protection of the host from SARS-CoV-2 infection.
